# Predicting Future Development of Stress-Induced Anhedonia From Cortical Dynamics and Facial Expression

**DOI:** 10.1101/2024.12.18.629202

**Authors:** Austin A. Coley, Kanha Batra, Jeremy Delahanty, Laurel R. Keyes, Rachelle Pamintuan, Assaf Ramot, Jim Hagemann, Christopher R. Lee, Vivian Liu, Harini Adivikolanu, Jianna Cressy, Caroline Jia, Francesca Massa, Deryn LeDuke, Moumen Gabir, Bra’a Durubeh, Lexe Linderhof, Reesha R. Patel, Romy Wichmann, Hao Li, Kyle B. Fischer, Talmo D. Pereira, Kay Tye

## Abstract

The current state of mental health treatment for individuals diagnosed with major depressive disorder leaves billions of individuals with first-line therapies that are ineffective or burdened with undesirable side effects. One major obstacle is that distinct pathologies may currently be diagnosed as the same disease and prescribed the same treatments. The key to developing antidepressants with ubiquitous efficacy is to first identify a strategy to differentiate between heterogeneous conditions. Major depression is characterized by hallmark features such as anhedonia and a loss of motivation (1, 2), and it has been recognized that even among inbred mice raised under identical housing conditions, we observe heterogeneity in their susceptibility and resilience to stress (3). Anhedonia, a condition identified in multiple neuropsychiatric disorders, is described as the inability to experience pleasure and is linked to anomalous medial prefrontal cortex (mPFC) activity (4). The mPFC is responsible for higher order functions (5–8), such as valence encoding; however, it remains unknown how mPFC valence-specific neuronal population activity is affected during anhedonic conditions. To test this, we implemented the unpredictable chronic mild stress (CMS) protocol (9–11) in mice and examined hedonic behaviors following stress and ketamine treatment. We used unsupervised clustering to delineate individual variability in hedonic behavior in response to stress. We then performed in vivo 2-photon calcium imaging to longitudinally track mPFC valence-specific neuronal population dynamics during a Pavlovian discrimination task. Chronic mild stress mice exhibited a blunted effect in the ratio of mPFC neural population responses to rewards relative to punishments after stress that rebounds following ketamine treatment. Also, a linear classifier revealed that we can decode susceptibility to chronic mild stress based on mPFC valence-encoding properties prior to stress-exposure and behavioral expression of susceptibility. Lastly, we used a markerless pose tracking computer vision tool, SLEAP (31), to predict whether a mouse would become resilient or susceptible based on facial expressions during a Pavlovian discrimination task. These results indicate that mPFC valence encoding properties and behavior are predictive of anhedonic states. Altogether, these experiments point to the need for increased granularity in the measurement of both behavior and neural activity, as these factors can predict the predisposition to stress-induced anhedonia.

## Introduction

Anhedonia—described as the inability to experience pleasure and hedonic feeling (12, 13) - is an underlying condition and core feature observed in both schizophrenia (SCZ), major depressive disorder (MDD)(14), and bipolar disorder (BD) (15, 16), and is suggested to be linked to anomalous medial prefrontal cortex (mPFC) activity (4). The mPFC, a higher order cortical region primarily responsible for cognition (5, 6), working memory (7, 8), sociability (17), and emotional control (18), is also involved in valence encoding (19), essential for discerning positive and negative hedonic values (20). Stress plays a major role in disrupting mPFC processes leading to depressive-phenotypes and is highly responsive to treatment. Ketamine administration shows promise as an antidepressant for treatment-resistant patients and has notable effects on mPFC cortical neurons (21–23). Indeed, mPFC imaging studies in MDD patients have identified biomarkers that can predict the response to therapy (24, 25). Recently, non-invasive approaches such as facial expression analysis have been used to capture the emotional state of a subject (26, 27). This led us to hypothesize that mPFC valence-encoding processes and behavioral features, including facial expression, can predict future stress-induced phenotypes and response to ketamine.

### Anhedonia classification predicts associative learning performance

To test this, we implemented the unpredictable chronic mild stress (CMS) protocol (9–11) (Fig. 1a, b) to induce anhedonia and assessed consummatory pleasure, despair, motivation, and sociability across weeks. We used sucrose preference test (SPT) as a measure of anhedonia (9,10) and utilized unsupervised k-means clustering to classify subjects into resilient and susceptible clusters (Fig. 1c-e). We then evaluated SPT scores in non-stressed (control), resilient, and susceptible mice. Our results showed susceptible mice display a significant reduction in sucrose preference following post-stress (Fig. 1f). However, we observed no differences in sucrose preference scores between non-stressed and stressed groups at the baseline, ketamine, and post-ketamine time points (Fig. 1f, g).

**Fig. 1.**
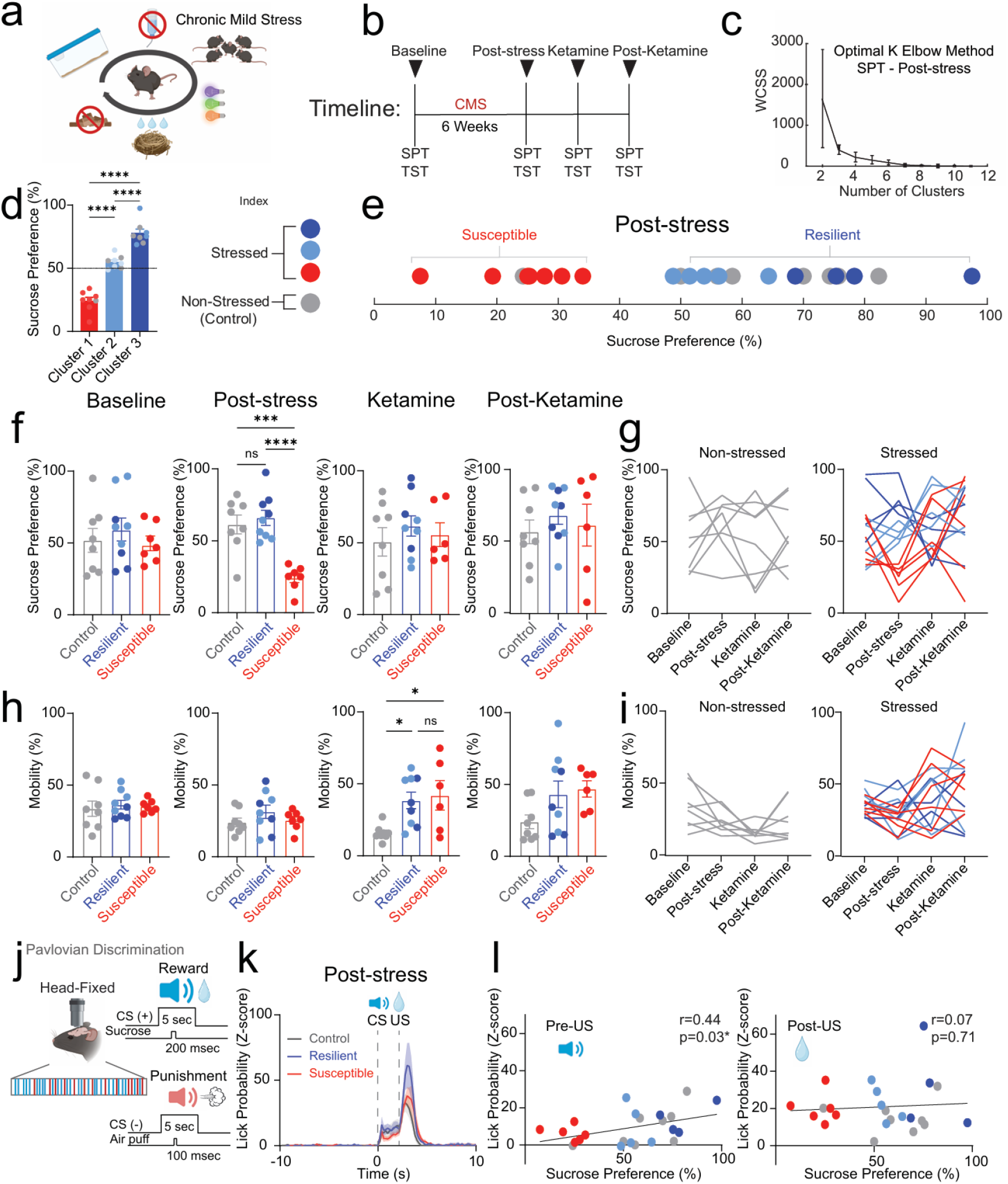
Stress-induced phenotype classification predicts reward task performance. **a**. Schematic of unpredictable chronic mild stress (CMS) protocol. CMS mice were exposed to 2-3 stressors per day for 6 weeks that consisted of cage tilting, strobe light illumination, white noise, crowded housing, light/dark cycle manipulations, food deprivation, water deprivation, and damp bedding. **b**. Timeline of measurements for sucrose preference test (SPT) and tail suspension test (TST) during CMS and ketamine treatment. **c**. The optimal k elbow method uses the within-cluster-sum-of-square (WCSS) values to determine the appropriate number of clusters derived from SPT scores of mice at the Post-stress time point. **d**. Cluster analysis of SPT scores for mice grouped in cluster 1, cluster 2, and cluster 3 at the post-stress time point. Significant decrease in SPT scores from cluster 1 mice compared to cluster 2 mice (One-way ANOVA, between-subjects F_(2,21)_=100.3, p<0.0001. Tukey post-hoc, p<0.0001). Significant decrease in SPT scores from cluster 2 mice compared to cluster 3 mice (p<0.0001). Significant increase in SPT scores from cluster 3 mice compared to cluster 1 mice (p<0.0001). **e**. To determine resilient (dark blue and light blue), and susceptible (red) subjects, k-means clustering (k=3) of sucrose preference scores was applied in both stressed (n=8) and non-stressed control (gray) groups (n=14). **f**. Susceptible mice displayed a reduction in SPT scores compared to control and resilient mice at the post-stress time point (One-way ANOVA, F_(2,21)_=16.95, p<0.0001, Tukey post-hoc: control compared to resilient mice, p=0.8051, control compared to susceptible mice, p=0.0003, susceptible compared to resilient mice, p<0.0001). No differences were observed at baseline (One-way ANOVA, F_(2,21)_=0.4606, p=0.6371), ketamine (One-way ANOVA, F_(2,20)_=0.4637, p=0.6356) or post-ketamine time points (One-way ANOVA, F_(2,20)_=0.4364, p=0.6524). **g**. Longitudinal description showing non-stressed control mice (left) and stressed (right) mice during sucrose preference test. **h**. Susceptible and resilient mice displayed an increase in mobility compared to control mice during TST at the ketamine time point (One-way ANOVA, F_(2,20)_=5.376, p=0.0135, Tukey post-hoc: control compared to resilient mice, p=0.0309; control compared to susceptible mice, p=0.0246; resilient compared to susceptible mice, p=0.9187. No differences in mobility across groups during baseline (One-way ANOVA, F_(2,21)_=0.3632, p=0.6997), post-stress (One-way ANOVA, F_(2,21)_=1.185, p=0.3253), and post-ketamine (One-way ANOVA, F_(2,20)_=2.702, p=0.0915) time points. **i**. Longitudinal description showing non-stressed control mice (left) and stressed (right) mice during tail suspension test. **j**. Pavlovian discrimination paradigm in a head-fixed mouse showing US paired with a 5-second pure tone as the conditioned stimulus (CS (+)), with the tone frequency set at 9 kHz for the rewarding CS (sucrose), and a 5-second pure tone as the conditioned stimulus (CS (-)), with the tone frequency set at 2 kHz for the punishment CS (air puff). **k**. Distribution of lick probability during reward trials in control, resilient, and susceptible mice. **l**. Significant correlation of lick probability and sucrose preference test during CS at Post-stress time point (Pearson’s correlation of lick probability and sucrose preference test in control, resilient, and susceptible mice. left, Pre-US, r=0.44, p=0.03; right, Post-US, r=0.07, p=0.71). Data in bar graphs are shown as mean and error bars around the mean indicate s.e.m. NS, not significant Error bars indicate s.e.m.

Additionally, CMS mice revealed no difference in mobility during tail suspension test (TST) at baseline or poststress time points, indicating no difference in behavioral despair or motivation; but showed an increase following ketamine treatment (Fig. 1h, i). These data suggest ketamine application reduces behavioral despair in stressed groups compared to control mice. We observed no significant differences in mobility across groups at the post-ketamine time point. Interestingly, we detected no difference in social preference in susceptible mice in response to CMS (Extended Data Fig. 1).

To assess the impact of chronic stress on the neural and behavioral readouts for reward or punishment-predictive cues, we trained mice that would ultimately undergo CMS or their non-stressed controls in a head-fixed Pavlovian discrimination task used to discriminate reward-predictive and punishment-predictive stimuli (Fig. 1j). During the task, one conditioned stimulus (CS) is paired (tone) with a 30% sucrose solution delivery reward (US-unconditioned stimulus), and a different CS is paired with a punishing air puff. We observed no significant differences in licking during the anticipatory phase (following CS onset and prior to US delivery) between stressed groups during the training phase (Extended Data Fig. 2). Our results showed no difference between groups in lick probability in reward trials during the anticipatory phase and consummatory phase (following US delivery) at the post-stress time point (Fig. 1k). Additionally, we measured lick probability during baseline, ketamine, and post-ketamine time points and observed no differences between groups during the CS or US phases (Extended Data Fig. 3). However, we did detect a significant correlation in lick probability and sucrose preference in all mice during the conditioned stimulus at the post-stress time point; suggesting that susceptible mice display both a reduction in lick probability and sucrose preference (Fig. 1l). No detectable correlation was observed during the unconditioned stimulus (Fig. 1l). These findings suggest that anhedonia classification can predict reward consumption performance during post-stress time points.

### Chronic stress blunts mPFC valence population dynamics and recovers at post-ketamine time point

To examine the relative dynamics of responses to reward-and punishment-predictive cues, we recorded longitudinal in vivo 2-Photon calcium imaging to track mPFC neuronal population activity (Extended Data Fig. 4), while mice are performing a Pavlovian discrimination task across 10 weeks during chronic mild stress and ketamine treatment (Fig. 2a-c). Using a local z-score (normalized to the baseline for each trial), we applied principal component analysis (PCA) to plot activity in a lower dimensional space during reward and punishment trials (Extended Data Fig. 5a). We examined population dynamics across weeks in non-stressed control, resilient and susceptible groups by measuring trajectory length post CS onset (0-10 sec) during reward trials and punishment trials (Extended Data Fig. 5b, c). Longer trajectories reflect more dynamic population activity during the trial (28). Our results showed no differences across groups during reward trials (Extended Data Fig. 5b). During punishment trials, we observed no differences in trajectory lengths at stress time points (Extended Data Fig. 5c).

**Fig. 2.**
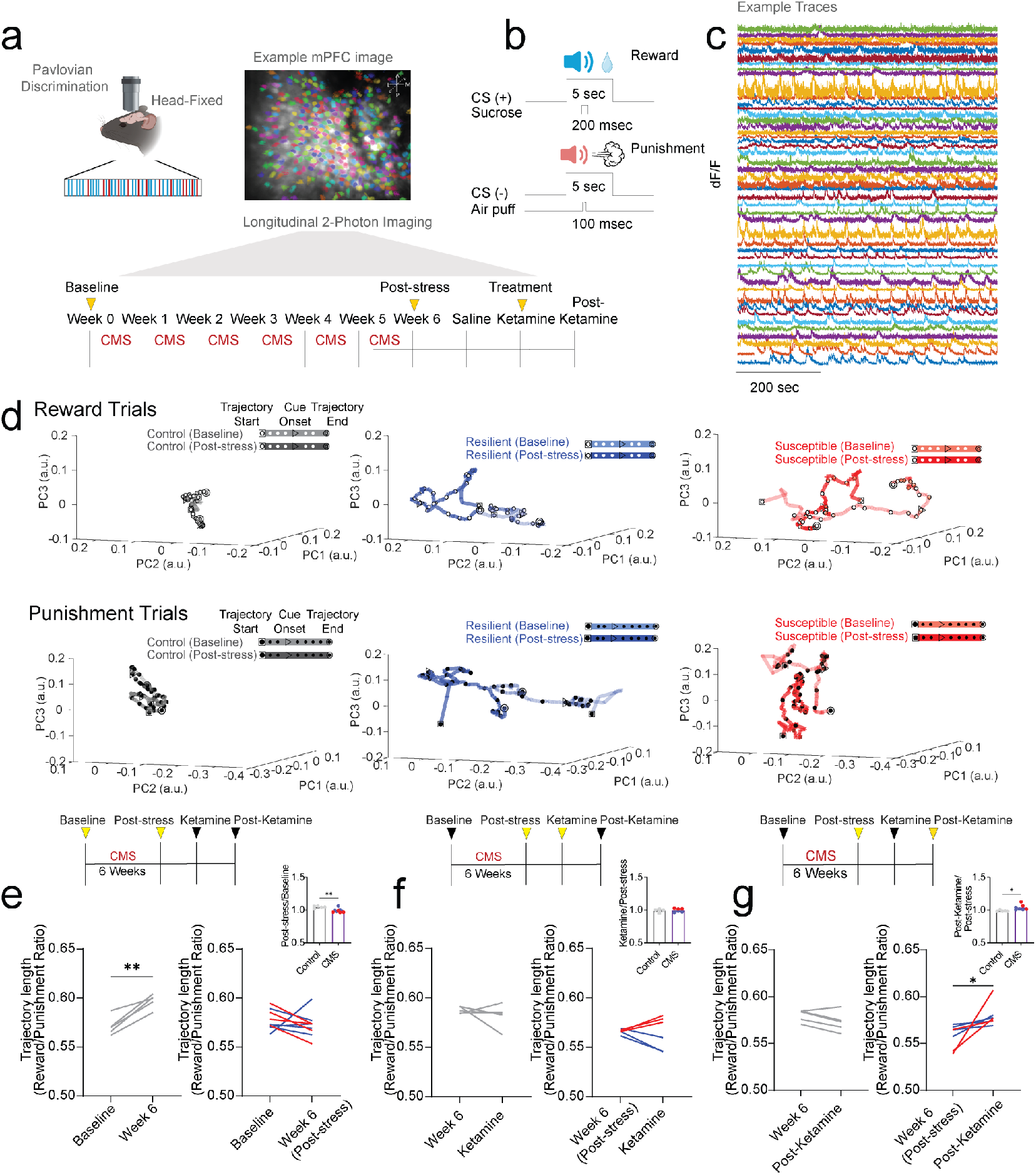
Chronic stress blunts mPFC valence population dynamics ratio while a single dose of ketamine reverses this effect. **a**. Head-fixed mouse and example mPFC 2-Photon image highlighting region of interest (ROI) neurons. Experimental paradigm shows the timeline of longitudinal 2-Photon imaging sessions. **b**. Pavlovian discrimination paradigm task showing Sucrose Reward trials (US paired with a 5-second tone (CS+)) and Air puff Punishment trials (US paired with a 5-second tone (CS-)). **c**. Example df/f traces of mPFC neurons. **d**. To explore population dynamics, we applied principal component analysis (PCA) of neural trajectories of ROI matched (co-registered) mPFC neurons during reward trials (Top) and punishment trials (Bottom) showing control (gray), resilient (blue), and susceptible (red) groups in a lower dimensional common principal component (PC) sub-space from Baseline to Post-stress time points. The first PCs capture 42.97% of the variance. The top 23 PCs were used to capture 59.51% of the variance. **e**. To examine the reward and punishment population dynamics we examined we used a super global Z-score (Z-score normalized across multiple sessions) and measured the trajectory lengths (post-event, 0-10 sec) during reward and punishment trials in pairwise (time point matched) ROI matched co-registered neurons and calculated the reward/punishment ratio during baseline to post-stress time points. Control mice showed an increase in reward/punishment ratio over time; Control (left), paired t-test, p=0.0031. Stressed mice showed no difference: CMS (right), paired t-test, p=0.3805. Significant decrease in trajectory length ratio (ratio normalized to baseline time point) in CMS mice compared to control mice. Bar graph: unpaired t-test, p=0.0031. **f**. No significant differences were observed in pairwise ROI matched neural trajectory lengths (post-event, 0-10 sec) reward/punishment ratio during Post-stress to Ketamine time points: Control (left), paired t-test, p=0.4520; CMS (right), paired t-test, p=0.8203. Bar graph: unpaired t-test, p=0.6929. **g**. Stressed groups showed an increase in reward/punishment ratio in pairwise ROI matched neural trajectory lengths (post-event, 0-10 sec) reward/punishment ratio during Post-stress to Post-Ketamine time points CMS (right), paired t-test, p=0.0475. No significant differences were observed in control groups. Control (left), paired t-test, p=0.0774. Significant increase in trajectory length ratio (ratio normalized to Post-stress time point) in CMS mice compared to control mice. Bar graph: unpaired t-test, p=0.0277. *p<0.05, **p<0.01.

To further evaluate the evolution of responses to reward-and punishment-predictive cues in mPFC neurons, we tracked and matched individual single cells over weeks and calculated the reward/punishment ratio of the PCA trajectory lengths in response to chronic stress and ketamine treatment as a reflection of the relative change in population dynamics (Fig. 2d; Extended Data Fig. 6a, b). Our results showed an increase in the reward/punishment ratio from baseline to week 6 (post-stress time point) in control mice, indicating an increase in mPFC reward processing over time (Fig. 2e). Subjects exposed to chronic mild stress displayed no difference in population dynamics ratio from baseline to poststress (Fig. 2e). We then measured the reward/punishment balance from post-stress to ketamine periods, and observed no difference in control or stressed groups (Fig. 2f, Extended Data Fig. 6a). Interestingly, when examining the difference from post-stress to post-ketamine time points we revealed an increase in the reward/punishment ratio of the PCA trajectory lengths in stress subjects (Fig. 2g, Extended Data Fig. 6b), indicating an increase in reward processing preference in both resilient and susceptible groups one week following ketamine treatment. We observed no difference in reward/punishment balance in control mice at post-stress to post-ketamine periods, suggesting stress-dependent changes in response to ketamine (Extended Data Fig. 6b).

### mPFC population activity predicts anhedonia phenotypes prior to stress exposure

To determine if mPFC population activity encodes stress-induced anhedonia behavioral phenotype classification, we used a generalized linear model (GLM) to predict if mPFC neuronal population activity could decode control, resilient and susceptible groups (Fig. 3a). We trained and tested neural data acquired from the first sucrose lick during reward trials and air puff during punishment-US across weeks, and analyzed decoding performance for resilient vs. control groups, susceptible vs. control groups, and resilient vs. susceptible groups. Our results showed there is a high decoding performance for resilient vs. control groups compared to shuffled data during first sucrose lick during individual weeks (Extended Data Fig. 7a). In susceptible vs. control groups, we observed a significantly greater decoding performance during sucrose lick at all time points; and most weeks were distinguishable for resilient vs. susceptible performance with the exception of week 1 (Extended Data Fig. 7b, c.). These data suggest that mPFC population activity can be used to discern susceptible and resilient phenotypes in response to first sucrose lick.

**Fig. 3.**
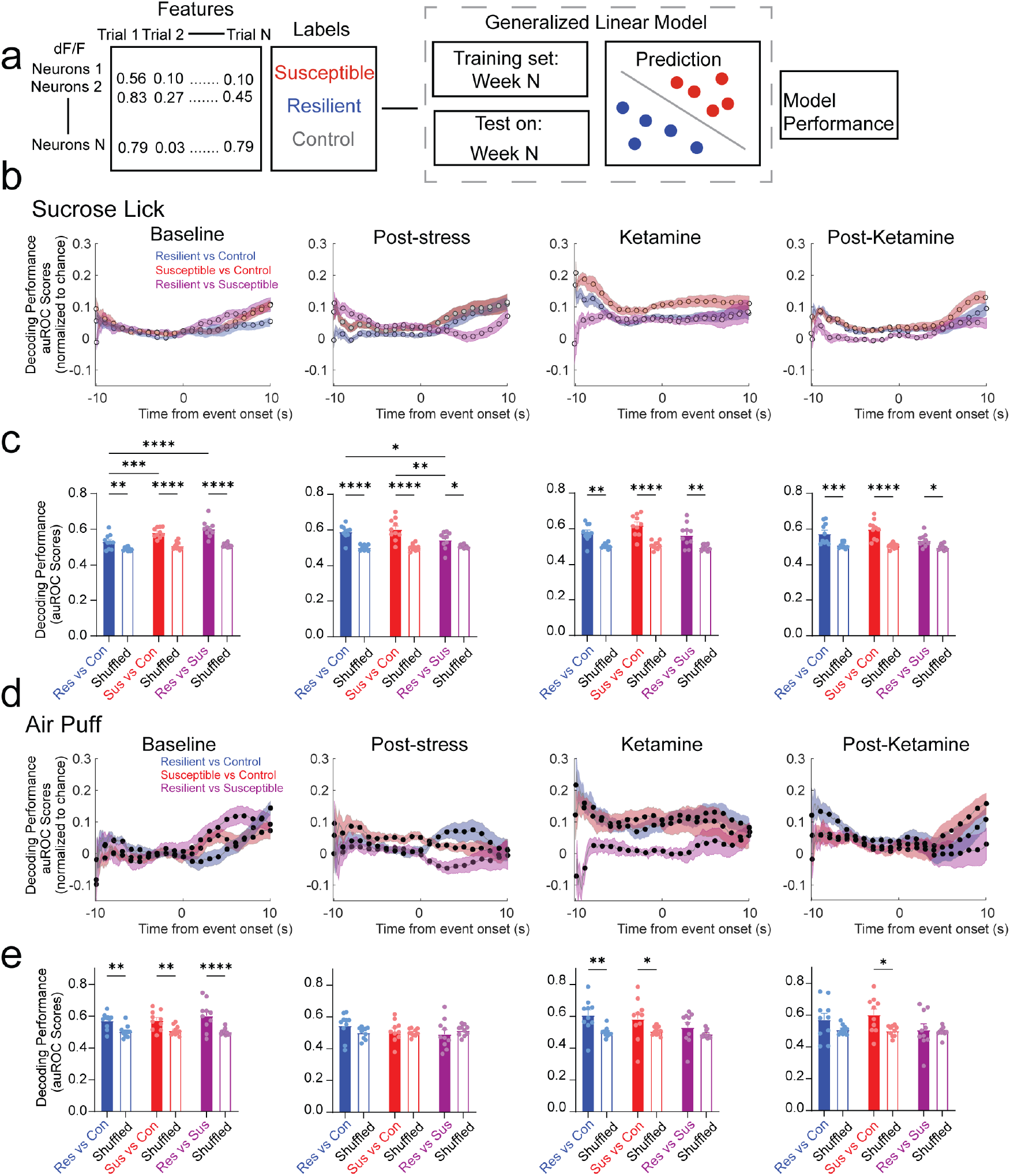
mPFC population dynamics predicts future resilience or susceptibility, before stress exposure. **a**. Schematic depicts feature and label inputs for the generalized linear model classifier used for decoding performance. **b**. Decoding performance across time during the first sucrose lick following US presentation (reward trials) in *resilient* vs. *control* groups (blue), *susceptible* vs. *control* groups (red), and *resilient* vs. *susceptible* groups (purple) at Baseline, Post-stress, Ketamine, and Post-Ketamine time points. **c**. Decoding performance during Sucrose lick (first lick following sucrose presentation). *Susceptible* vs. *control* groups displayed a significantly greater decoding performance than *resilient* vs. *control* groups at Baseline (Two-way ANOVA, event F_(1,27)_=86.98, p<0.0001, groups F_(2,27)_=11.91, p=0.0002, interaction, F_(2,27)_=4.175, p=0.0263; Tukey post-hoc, *resilient* vs. *control* compared to *susceptible* vs. *control* groups, p=0.0010, *resilient* vs. *control* compared *resilient* vs. *susceptible* groups. p<0.0001). Significantly greater decoding performance in *resilient* vs. *control* compared to *resilient* vs. *susceptible* groups, and *susceptible* vs. *control* groups compared to *resilient* vs. *susceptible* groups at the Post-stress time point (Two-way ANOVA, event F_(1,27)_=58.08, p<0.0001, groups F_(2,27)_=3.0009, p=0.0661, interaction, F_(2,27)_=4.110, p=0.0277; Tukey post-hoc, *resilient* vs. *control* compared to *resilient* vs. *susceptible* groups, p=0.0216, *susceptible* vs. *control* groups compared to *resilient* vs. *susceptible* groups, p=0.0019). Stress phenotypes displayed a significantly higher decoding performance compared to shuffled data at each time point, but no differences were observed across groups at Ketamine (Two-way ANOVA, event F_(1,27)_=203.4, p<0.0001, groups F_(2,27)_=3.693, p=0.0382, interaction, F_(2,27)_=1.450, p=0.2522) and Post-Ketamine time points (Two-way ANOVA, event F_(1,27)_=55.42, p<0.0001, groups F_(2,27)_=4.134, p=0.0272, interaction, F_(2,27)_=3.203, p=0.0564). **d**. Time series traces depicting decoding performance during air puff-US (punishment trials) in *resilient* vs. *control* groups (blue), *susceptible* vs. *susceptible* groups (red), and *resilient* vs. *susceptible* groups (purple) at Baseline, Post-stress, Ketamine, and Post-Ketamine time points. **e**. Decoding performance during air puff-US. Significantly greater decoding performance of *resilient* vs. *control* groups and *susceptible* vs. *control* groups compared to shuffled data at Baseline (Two-way ANOVA, event F_(1,27)_=41.80, p<0.0001, groups F_(2,27)_=0.2737, p=0.7627, interaction, F_(2,27)_=1.056, p=0.3617), and the Ketamine time points (Two-way ANOVA, event F_(1,27)_=14.46, p=0.0007, groups F_(2,27)_=1.437, p=0.2552, interaction, F_(2,27)_=0.9261, p=0.4083), but no differences across stress groups. No difference in decoding performance of *resilient* vs. *susceptible* groups to shuffled data and *susceptible* vs. *susceptible* groups compared to shuffled data at the Post-stress time point (Two-way ANOVA, event F_(1,27)_=0.3822, p=0.5416, groups F_(2,27)_=0.6345, p=0.5379, interaction, F_(2,27)_=1.679, p=0.2054). Mice with a susceptible phenotype displayed a significantly greater decoding performance compared to shuffled data at Post-Ketamine time point, but no differences were observed across stress groups (Two-way ANOVA, event F_(1,27)_=65.09, p<0.0001, groups F_(2,27)_=1.840, p=0.1782, interaction, F_(2,27)_=1.392, p=0.2659). All post-hoc comparisons are Tukey t-tests, *p<0.05, **p<0.01, ***p<0.001, ****p<0.0001. All 2-way ANOVAs were for event (event vs. shuffle) and groups (*resilient* vs. *susceptible, susceptible* vs. *susceptible*, and *resilient* vs. *susceptible*). Data in bar graphs are shown as mean and error bars around the mean indicate s.e.m.

We then compared decoding performance between stress groups at baseline, post-stress, ketamine, and postketamine time points during first sucrose lick. Interestingly, at baseline, we observed a significant increase in decoding performance in susceptible vs. control groups compared to resilient vs. control groups (Fig. 3b, c). These data suggest that mPFC neural population activity in susceptible mice is more distinct compared to resilient mice in response to reward stimuli prior to stress. Additionally, at the post-stress, ketamine, and post-ketamine time points, we observed a significantly greater decoding performance in both susceptible vs. control and resilient vs. control groups compared to the resilient vs. susceptible group. These data indicate mPFC population activity can decode anhedonia phenotypes during stress and ketamine treatment in response to first sucrose lick.

Next, we examined decoding performance in response to air puff between resilient vs. control groups, susceptible vs. control groups, and resilient vs. susceptible groups across weeks (Extended Data Fig. 7d-f). The susceptible vs. control groups displayed a significant increase in decoding performance compared to shuffle data within individual weeks except at an early stress time point (week 2), and late stress time points (weeks 4-6), and saline and ketamine administration sessions (weeks 7-8) (Extended Data Fig. 7e). Interestingly, we observed no significant differences in decoding performance across weeks in resilient vs. control groups or resilient vs. susceptible groups in response to air puff stimuli (Extended Data Fig. 7d, f). These data suggest that susceptible vs. control groups display distinct mPFC activity encoding properties in response to air puff during stress.

To measure the difference in resilient vs. control, susceptible vs. control, and resilient vs. susceptible groups in response to air puff stimuli we measured the decoding performance at baseline, post-stress, ketamine, and post-ketamine time points (Fig. 3d). At baseline, we were able to significantly decode resilient mice from control mice, susceptible from control, and resilient from susceptible groups compared to shuffled data (Fig. 3e). But, unlike in reward trials, we did not detect a difference amongst the resilient vs. control compared to susceptible vs. control at baseline for punishment trials (Fig. 3e). During post-stress, the resilient vs. control, susceptible vs. control, and resilient vs. susceptible groups displayed no difference compared to shuffled data (Fig. 3e). These data demonstrate chronic mild stress ablates phenotype decoding performance during punishment trials.

### Facial expression features track changes from baseline and decode future stress phenotypes

To evaluate the affective state of subjects exposed to chronic stress, we used markerless pose tracking system SLEAP (31) to examine the facial features in response to reward and punishment trials (Fig. 4a). To capture the spatiotemporal dynamics of facial expressions, we extracted high dimensional facial data from videos during the head-fixed Pavlovian discrimination task and then plotted these features in reduced dimensional space using principal component analysis to track facial expression dynamics before, during, and after stress, as well as after ketamine treatment (Fig. 4b). Similar to neural analysis, using a local z-score (10 sec prior to CS-onset), we examined facial dynamics prior to stress exposure in control, resilient, and susceptible groups by measuring the difference score (post-event - baseline) of facial trajectory lengths during reward trials (Supplementary Video 1). At baseline, our results show that the difference score between facial trajectory lengths during the anticipatory window (CS-onset to 2 sec post CS-onset) prior to reward trials is decreased in susceptible mice compared to resilient mice suggesting that facial expression dynamics can predict future behavioral phenotypes induced by stress (Fig. 4c). There was no increase in licking during anticipatory periods during the baseline, therefore the difference in facial expression between resilient and susceptible mice was not due to differences in mouth movement due to licking (Extended Data Figure 8).

**Fig. 4.**
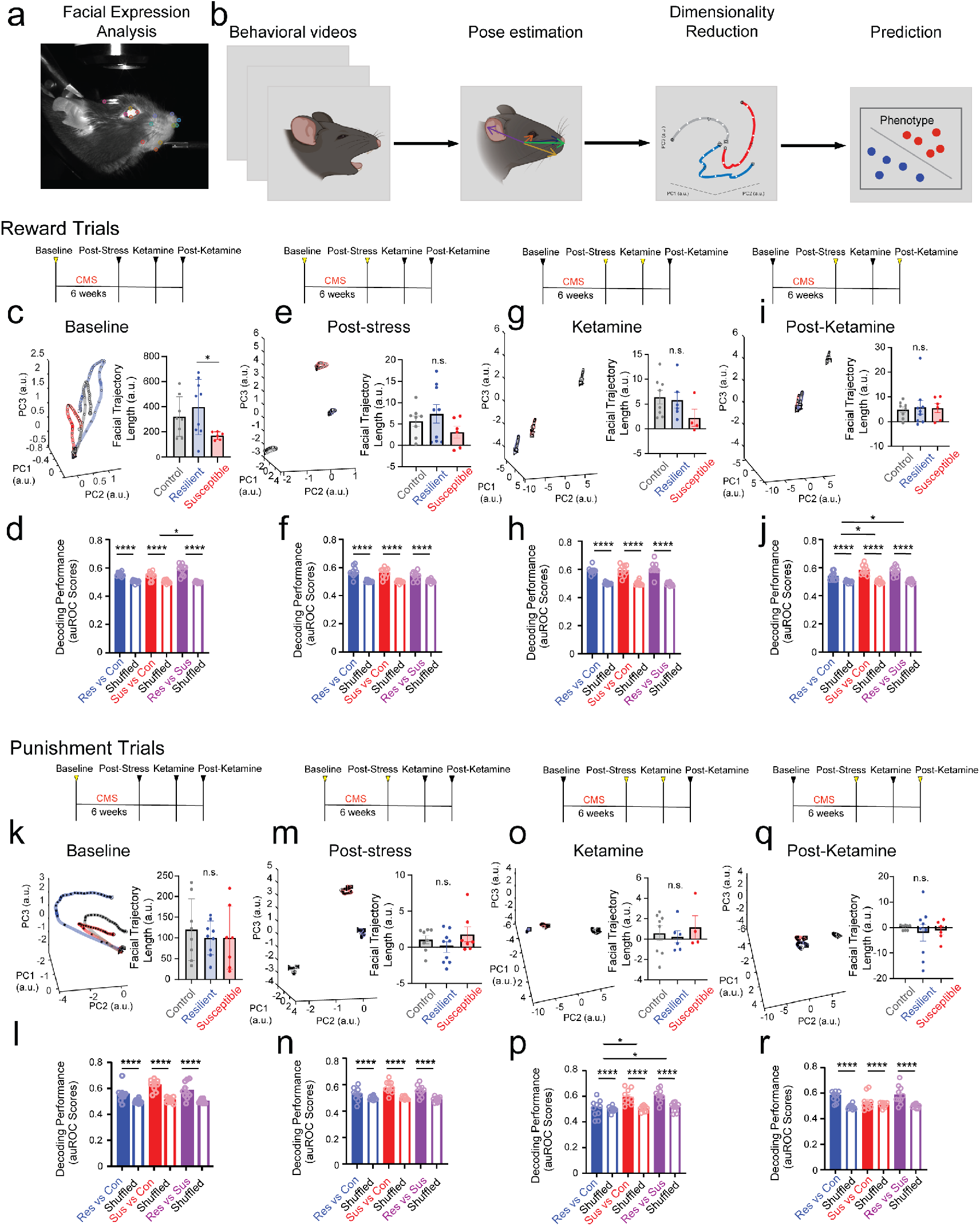
Susceptibility and resilience can be robustly decoded and predicted from facial expression dynamics. **a**. Example image of labeled mouse facial features. **b**. To determine if we could predict future responses to stress or responses to ketamine based on facial features alone, we first extracted facial keypoints from using SLEAP, then plotted the facial expression dynamics in a dimensionality-reduced trajectory in time across principal component space of facial expression dynamics. **c**. To measure facial dynamics, we used a local Z-score, and extracted PCA trajectories (top 3 PCs capture 81.91% of the variance; 8 PCs were used to capture 90.54% of the variance) of facial features at baseline (left) and difference score (right) of trajectory lengths post-event (10 sec CS) – pre-event (10 sec Pre-CS). Control and resilient groups displayed a significantly greater PCA difference score compared to susceptible mice. One-way ANOVA, between-subjects F_(2,21)_=20.18, p<0.0001. Tukey post-hoc, control compared to resilient mice, p=0.9994, control compared to susceptible mice, p<0.0001, resilient compared to susceptible mice, p<0.0001) **d**. (FIX THIS SENTENCE) Significantly greater facial decoding performance in stressed groups compared to shuffled data, but no difference across stressed groups during reward trials at Baseline (Two-way ANOVA, event F_(1,18)_=573.2, p<0.0001, groups F_(2,36)_=3.095, p=0.0575, interaction, F_(2,36)_=3.206, p=0.0523). **e**. Susceptible groups displayed a significant increase in PCA difference score compared to control and resilient groups at post-stress (One-way ANOVA, F_(2,21)_=9.139, p=0.0014. Tukey post-hoc, control compared to resilient mice, p=0.9784, control compared to susceptible mice, p=0.0045, resilient compared to susceptible mice, p=0.0023). **f**. Significantly greater facial decoding accuracy in stressed groups compared to shuffled data, but no difference across groups at Post-stress time point. (Two-way ANOVA, event F_(1,18)_=344.3, p<0.0001, groups F_(2,36)_=0.1186, p=0.8885, interaction, F_(2,36)_=1.898, p=0.1645). **g**. Resilient mice displayed a significant reduction in PCA difference score compared to control and susceptible groups following ketamine administration (One-way ANOVA, F_(2,15)_=15.18, p=0.0002. Tukey post-hoc, control compared to resilient mice, p=0.0002, control compared to susceptible mice, p=0.3646, resilient compared to susceptible mice, p=0.0143). **h**. Significantly greater decoding performance in stressed groups compared to shuffled data, and a significantly greater increase in *susceptible vs. control* groups compared to *resilient vs. control* groups at ketamine time point (Two-way ANOVA, event F_(1,18)_=255.9, p<0.0001, groups F_(2,36)_=21.30, p<0.0001, interaction, F_(2,36)_=5.525, p=0.0081. Tukey post-hoc, *resilient vs. control* resilient compared to *susceptible vs. control* groups, p<0.0001, *resilient vs. control* groups compared to *resilient vs. susceptible* groups, p=0.0068, *susceptible vs. control* groups compared to *resilient vs. susceptible* groups, p=0.0023. **i**. Resilient mice display a significant reduction in PCA difference score compared to control and susceptible groups at post-ketamine (One-way ANOVA, F_(2,20)_=9.206, p=0.0015. Tukey post-hoc, control compared to resilient mice, p=0.0054, control compared to susceptible mice, p=0.9070, resilient compared to susceptible mice, p=0.0038. **j**. We found a significantly greater decoding performance in stressed groups compared to shuffled data, and a significantly higher decoding performance in *resilient vs. control* groups compared to *resilient vs. susceptible* groups and *susceptible vs. control* compared to *resilient vs. susceptible* groups at the post-ketamine time point (Two-way ANOVA, event F_(1,18)_=665.3, p<0.0001, groups F_(2,36)_=6.825, p=0.0031, interaction, F_(2,36)_=5.316, p=0.0095. Tukey post-hoc, *resilient vs. control* compared to *susceptible vs. control*, p=0.6321, *resilient vs. control* groups compared to *resilient vs. susceptible* groups, p=0.0019, *susceptible vs. control* groups compared to *resilient vs. susceptible* groups, p=0.0001). **k**. Resilient groups displayed a significant increase in PCA difference score compared to control and susceptible groups at Baseline during punishment trials (One-way ANOVA, F_(2,21)_=10.85, p=0.0006. Tukey post-hoc, control compared to resilient mice, p=0.0016, control compared to susceptible mice, p>0.9999, resilient compared to susceptible mice, p=0.0023). **l**. We observed a significantly greater decoding performance in stressed groups compared to data with shuffled labels, and a significantly greater increase in *resilient vs. control* compared to *susceptible vs. control* groups at baseline (Two-way ANOVA, event F_(1,18)_=91.33, p<0.0001, groups F_(2,36)_=7.033, p=0.0026, interaction, F_(2,36)_=4.068, p=0.0255. Tukey post-hoc, *resilient vs. control* groups compared to *susceptible vs. control* groups, p=0.0077, *resilient vs. control* groups compared to *resilient vs. susceptible* groups, p=0.3890, *susceptible vs. control* compared to *resilient vs. susceptible* groups, p=0.0002). **m**. No differences in PCA difference scores at the post-stress time point (One-way ANOVA, F_(2,21)_=2.884, p=0.0782). **n**. Significantly greater decoding performance in stressed groups compared to shuffled data, but no difference across groups at post-stress session (Two-way ANOVA, event F_(1,18)_=230.7, p<0.0001, groups F_(2,36)_=3.343, p=0.0466, interaction, F_(2,36)_=2.133, p=0.1357). **o**. Resilient mice displayed a significant reduction in PCA difference score compared to control and susceptible groups at ketamine time point (One-way ANOVA, F_(2,15)_=4.651, p=0.0268. Tukey post-hoc, control compared to resilient mice, p=0.0256, control compared to susceptible mice, p=0.9246, resilient compared to susceptible mice, p=0.1224). **p**. Significantly greater decoding performance in stressed groups compared to shuffled labels following ketamine administration, but no difference across groups (Two-way ANOVA, event F_(1,18)_=70.99, p<0.0001, groups F_(2,36)_=0.02305, p=0.9772, interaction, F_(2,36)_=1.060, p=0.3571). **q**. Susceptible mice displayed a significant increase in PCA difference score compared to control and resilient groups (One-way ANOVA, F_(2,20)_=12.58, p=0.0003. Tukey post-hoc, control compared to resilient mice, p=0.6814, control compared to susceptible mice, p=0.0022, resilient compared to susceptible mice, p=0.0003. **r**. Significantly greater decoding performance in stressed groups compared to shuffled data at post-ketamine, but no difference across groups (Two-way ANOVA, event F_(1,18)_=56.50, p<0.0001, groups F_(2,36)_=0.2553, p=0.7915, interaction, F_(2,36)_=0.3098, p=0.7355). Data in bar graphs are shown as mean and error bars around the mean indicate s.e.m.

To monitor changes in facial dynamics during the stress paradigm, we normalized facial features to the subject baseline to track individual changes due to stress. We performed PCA using baseline data only, and projected data from subsequent weeks onto the resulting reduced dimensional space. We found no significant changes between groups during the anticipatory windows at post-stress (Fig. 4e). However, facial dynamics are readily trackable across weeks with a strong correlation between the first two principle components in the control and resilient groups. Interestingly, this linear relationship breaks down in susceptible mice where, across sessions, trajectories drift to the lower right quadrant of the PC space (Extended Data Figure 9a-c).

To quantify the strength of the correlation between PC1 and PC2, we examined the linear trend of the trajectories in principal component space. The alignment of trajectories along the diagonal of the PC1-PC2 place over time suggests that this relationship can be captured by focusing on the intertrial interval (ITI). We extracted facial features 10 sec before tone onset, then trial-averaged for each subject and each week. Over the 10 weeks, we found a strong correlation between PC1 and PC2 for control and resilient groups but no detectible correlation for susceptible mice (Extended Data Figure 9g-i).

To explore the interpretation of the principal components, we sorted the weights associated with each facial feature for PC1 and PC2 and found that the highest weighted features of PC1 appeared to track eye-related features, while for PC2, the highest weights related to mouth features (Fig. 10, 11). Based on this interpretation, we find that resilient and control animals have a strong correlation between mouth movements and eye-blinks but this is not true for susceptible animals.

To explore facial dynamics during ketamine and postketamine sessions, we normalized the facial features to the post-stress session, since we wanted to test whether ketamine could ameliorate the impact of CMS on facial expression dynamics. We performed PCA using the post-stress week only, and projected data from the ketamine and post-ketamine sessions onto the resulting reduced dimensional space. No significant changes were noted between groups during the anticipatory windows following ketamine administration or post-ketamine sessions (Fig. 4g, i).

We measured facial dynamics across weeks during stress and ketamine treatment in control, resilient, and susceptible groups by analyzing trajectory lengths post-event (CS-onset to 10 sec post-CS-onset) during reward trials (Extended Data Fig. 12a-c). Our results showed control mice exhibited a dramatic increase in facial dynamics during the reward tone immediately following ketamine administration, when normalized to their post-stress session (Extended Data Fig. 12a). Surprisingly, stressed mice neither resilient nor susceptible, showed changes to facial trajectory lengths across weeks (Extended Data Fig. 12b, c).

To test whether we could predict if facial responses to reward stimuli could decode control, resilient and susceptible groups, we applied a generalized linear model to trialaveraged facial features in the reduced dimensional space (Fig. 4d). We showed efficient decoding performance of stress groups for sucrose trials across weeks (Extended Data Fig. 13a-c). We observed a significant increase in facial decoding performance in stress phenotypes compared to data with shuffled labels at each of the time points considered, baseline, post-stress, ketamine, and post-ketamine (Fig. 4d, f, h, j). Interestingly, our results also showed a significantly higher decoding performance in resilient vs. susceptible groups compared to susceptible vs. control groups at baseline during reward trials (Fig. 4d). Furthermore, overall decoding performance was higher in both susceptible vs. control and susceptible vs. resilient groups compared to resilient vs. control (Fig 4j).

Next, we examined facial dynamics across weeks in control, resilient and susceptible groups by measuring the difference score (post-event - baseline 10 sec prior to CSonset) of trajectory lengths during punishment trials. As in analysis of reward trials above, we used a local z-score normalization (10 sec prior to CS-onset) for the baseline session but found no significant differences in the difference score of facial trajectory lengths during the anticipatory window at baseline (Fig. 4k).

To track changes due to stress from an unstressed baseline state, we normalized stress weeks to baseline; we performed PCA on the baseline data only, and projected subsequent weeks into the resulting reduced dimesional space. We found no significant difference between groups in the difference score of facial trajectory lengths during the anticipatory window at the post-stress session in response to punishment (Fig. 4m). However, as with the reward trials, there is a strong linear correlation between PC1 and PC2 in control and resilient animals during punishment trials across all weeks (Extended Data Fig 9d, e). This correlation is not detected for susceptible animals (Extended Data Fig 9f). Instead data from susceptible mice appears to drift over time to the lower right quandrant of the PC space similar to the pattern noted for susceptible mice during reward trials. This suggests that the correlation seen in resilient and control animals between eye-blinks and mouth movements are absent in susceptible animals (Extended Data Fig 9g-i, 10, 11).

To understand and track changes due to ketamine from the stressed state, we normalized session data collected following ketamine administration and the post-ketamine session (one week following ketamine) to the post-stress session. We performed PCA on the post-stress session only, and projected the ketamine and post-ketamine session data into the resulting reduced dimensional space. We found no significant difference between groups in the difference score of facial trajectory lengths during the anticipatory window following ketamine administration, nor during the post-ketamine session (Fig. 4o, q).

We then measured facial dynamics across weeks during stress and ketamine treatment in control, resilient, and susceptible groups during the response window for punishment trials (Extended Data Fig. 12d-f). Our results showed an increase in trajectory lengths in resilient mice during saline administration and during post-ketamine session (Extended Data Fig. 12e). No changes across weeks were identified for control or susceptible groups (Extended Data Fig. 12d, f).

To test facial decoding performance in response to air puff between stress groups, we used a GLM and showed efficient decoding performance across weeks in resilient vs. control during stress weeks and susceptible vs. control groups over all weeks (Extended Data Fig. 13d,e). Notably, the weeks for the stress paradigm were less decodable for the resilient vs. susceptible group during punishment trials indicating that these groups share similar facial responses to punishment during stress (Extended Data Fig. 13f).

Next, we compared decoding performance between resilient vs. control groups, susceptible vs. control groups, and resilient vs. susceptible groups during punishment trials at baseline, post-stress, ketamine, and post-ketamine time points (Fig. 4l, n, p, r). Following ketamine application, we noticed an increase in both resilient vs. susceptible and susceptible vs. control decoding performance compared to resilient vs. control (Fig. 4p). However, we observed no difference among groups at baseline, following post-stress or post-ketamine (Fig. 4l, n, r). These results indicate that ketamine appears to increase facial dynamics for susceptible mice during punishment stimuli; however, in other sessions, facial dynamics are not significantly different across groups in response to punishment.

## Conclusion

Together, these data reveal that mPFC valencespecific neural population activity and behavioral attributes predict anhedonia phenotypes. Our data demonstrate that longitudinal tracking of neural populations and activity across epochs of unpredictable chronic mild stress can help identify biomarkers for depressive-like phenotypes. We demonstrate that mPFC neural dynamics and facial expression features can encode anhedonia at multiple time points. Susceptible mice displayed significantly decreased facial dynamics during the anticipatory reward period compared to resilient mice and significantly higher reward decoding performance compared to resilient mice at baseline before animals undergo stress, suggesting we can predict susceptibility prior to stress using facial expression alone. Interestingly, chronic stress eliminates the neural decoding performance of punishment unconditioned stimuli in both resilient and susceptible groups.

We investigated the differential effects of ketamine application in both control and stressed groups, showing alleviation of anhedonia phenotypes after a 24 hour recovery period and found that allieviation was sustained one week later. However, we demonstrate ketamine’s distinct stress-dependent changes during despair assays, where control mice show a reduction in mobility compared to both resilient and susceptible groups. Our data also highlight a preference in mPFC reward processing in stressed groups one week after ketamine administration. These data support the decoding studies, showing that susceptible mice exhibit higher decoding performance compared to resilient mice, which we speculate reflects an increased sensitivity to ketamine application within PFC dynamics and associated facial feature expressions. These data could lead to ketamine response predictions and sustainability, poised for subjects exposed to chronic stress. Altogether, this study highlights the importance of longitudinal data as a framework for identifying biomarkers of depressive-like phenotypes by analyzing granular behavioral attributes in combination with mPFC neural dynamic population features.

## Acknowledgements

We thank Takaki Komiyama for technical support. Sotoris Masmanidis and Bitna Joo for helpful comments on the manuscript. Salk machine shop and Salk behavior core for mechanical assistance. This preprint used the zHenriquesLab template for Overleaf and was formatted in MikTex.

## Author Contributions

A.A.C and K.M.T. conceived the project, designed and supervised the experiments. A.A.C, J.D. and R.W. performed stereotaxic surgeries. A.A.C, J.D., R.P., V.L., H.A., J.C., C.J., F.M., M.G., and L.L. performed behavioral experiments. A.A.C, J.D., C.J., K.F. performed 2-Photon calcium-imaging experiments. A.A.C, K.B., J.D., A.R., R.P., J.H., F.M. and H.L. processed and analyzed calcium data. K.B., R.P, L.R.K, C.R.L, M.G, B.D., and A.E. performed SLEAP automated pose tracking analysis for facial and social experiments. L.R.K and K.B. SLEAP facial expression analysis. R.P. performed histological verifications. A.A.C, K.B., J.D., A.R., L.R.K., J.H., C.L., D.L., R.R.P., A.E., H.L., K.B. provided code scripts, edited code and offered advice for data analysis. T.D.P. made additional significant intellectual contributions. A.A.C, K.B., and L.R.K graphed data and made figures. A.A.C and K.M.T. wrote the paper.

## Data and Materials Availability

All experimental data are available in the main text or supplementary material. Data will be available on an open-source database upon publication.

## Code availability

The MATLAB toolbox Facial Expression Feature Extration used to extract facial features in this manuscript is available on the Tyelab GitHub (https://github.com/Tyelab/FEFE).

## Supplementary Materials

Materials and Methods, Extended Data Figs. 1 to 13, Supplementary Video S1

## MATERIALS AND METHODS

### Animals and housing

Adult, male HET DAT-Cre genotyped mice (at the minimum age of 8 weeks) arrived from Jackson Laboratory (RRID: IMSRJAX:000,664) and bred at the Salk Institute, were used for this study. The mice were housed in a reverse light cycle, with ad libitum access to food and water, until the commencement of major survival surgery, behavioral tests or imaging sessions. The animals were accommodated in cages with up to three littermates mates. All animal handling procedures adhered to the guidelines stipulated by the National Institute of Health (NIH) and were approved by the UCSD Institutional Animal Care and Use Committee (IACUC).

### Stereotaxic surgeries

Under aseptic conditions, surgery was conducted on all subjects using a small animal stereotax (David Kopf Instruments, Tujunga, CA, USA), with body temperature maintenance achieved using a heating pad. Anesthesia was induced using a 5% mixture of isoflurane and oxygen, which was subsequently reduced to 2–2.5% and maintained throughout the procedure (0.5 L/min oxygen flow rate). Once the subjects reached an adequate level of anesthesia, measured using a toe pinch, a 1mg/kg Buprenorphine-SR injection was administered subcutaneously, the ophthalmic ointment was applied to protect the eyes, hair was clipped from the incision site, the area was scrubbed alternatively three times with betadine and 70% ethanol, and lidocaine was subcutaneously (SQ) injected at the incision site. All measurements for viral injections were referenced from Bregma as the origin. Following the surgery, the subjects were IP injected with 1mL Ringer’s Lactate and placed in clean cages containing water-softened mouse chow to facilitate recovery. The cages were positioned on a heating pad to aid in the recovery process.

### Viral injection and GRIN lens placement surgery

To enable recordings from medial prefrontal cortex (mPFC) neurons, a viral approach was implemented. Following the aforementioned general surgical procedures, an incision was made to expose the skull. After skull leveling, craniotomies were performed above the mPFC regions. For expression of GCaMP, 300 nL of AAV1-hSyn-jGCaMP7f was injected into the mPFC at stereotaxic coordinates of 1.9 mm anteroposterior, 0.40 mm mediolateral, and-2.2 mm dorsoventral from Bregma. The injections were carried out using a 10 µL Nanofil syringe (WPI, Sarasota, FL, USA) driven at a rate of 0.1 µL/min with a microsyringe pump and controller (Micro4; WPI, Sarasota, FL, USA). Following each viral injection, the needle was allowed to stay in place for 5-10 minutes to allow viral material penetration before extraction. To prevent contamination, the needle was thoroughly flushed with 70% ethanol and sterile water. Viral aliquots were sourced from Addgene (Watertown, MA). Subsequent to viral injections, a 1 x 4 mm gradient refractive index (GRIN) lens (Proview, Inscopix Inc, Mountain View, CA, USA) was inserted into the mPFC at stereotaxic coordinates of 1.9 mm anteroposterior, 0.4 mm mediolateral, and-2.18 mm dorsoventral from Bregma. The GRIN lens was then secured to the skull and headplate using C&B Metabond and cement (Parkell), respectively.

### Behavioral testing

All behavioral testing occurred after a minimum of three weeks post-surgery recovery. Mice were individually handled for 15 minutes each day for five days to gain familiarity with experimenters and reduce stress during experiments.

### Sucrose preference test

The sucrose preference test (SPT) was used to measure anhedonia and was conducted in operant chambers (Med Associates, Inc) placed within sound-attenuated cubicles. Each SPT session lasted for 60 minutes and involved the use of two electrical lickometers and a house light set at an intensity of 40 lux. The lickometers were connected to bottles containing either tap water or a 1% sucrose solution in tap water. The MedPC IV software (Med Associates, Inc) was utilized to detect and record each lick event. Sucrose preference was calculated as (sucrose lick /(sucrose lick + water lick)) x 100. No additional food sources were available within the operant chambers. To ensure variability, the bottle configuration was different in each of the six operant chambers used. This allowed for repeated measures experiments, enabling animals to be re-tested and re-establish learning during each session.

### 3-Chamber Sociability test

The 3-chamber sociability test was used to measure sociability and was performed in a clear rectangular plexiglass arena. Prior to each session, the subject mouse was habituated in the empty arena for 3 minutes. Subsequently, the mouse was taken out of the arena, and a novel male mouse was placed inside a barred cup on one side of the arena together with an empty barred cup on the opposite side. The subject mouse was placed in the arena for 7 minutes during which footage was taken with a digital video camera above the arena. Ethovision XT software (Noldus, Wageningen, Netherlands) was used to record the mice during sociability assay.

### Tail suspension test

The tail suspension test was used to measure behavioral despair. The tail of each mouse was placed between two strips of autoclave labeling tape. The end of one strip of tape was then secured to a horizontal bar 40 cm from the ground, ensuring that the animal could not make other contact or climb during the assay. Video recording was started 90 s from the time that the animal was inverted and taped. Mice were inverted for 6 minutes. Time spent struggling was measured by OD-log and blind scoring each minute of video material after the testing was completed and was reported in seconds for each minute of the assay.

### Unpredictable Chronic Mild Stress protocol

To induce anhedonic symptoms, the chronic mild stress (CMS) protocol was implemented within a mouse model (11). Mice in the CMS group were exposed to 2-3 stressors per day for 6 weeks that consisted of cage tilting, strobe light illumination, white noise, crowded housing, light/dark cycle manipulations, food deprivation, water deprivation, and damp bedding. CMS mice were exposed to ∼3-4 hours per day besides the 12 hr light/dark cycle stressors. Stressors were imposed over all cages and randomized across all the days. Control mice were not exposed to stressors.

### Ketamine administration

After the 6-week chronic mild stress protocol, all mice were IP injected with saline (0.01-0.04 ml). The following week all mice were IP injected with ketamine (10 mg/kg, 0.01-0.04 ml) to alleviate anhedonia. Mice were allowed to recover at least 24 hours after injection before performing behavioral tasks or imaging experiments.

### Anhedonia Classification

Mice were classified following chronic mild stress using unsupervised k-means clustering method (k=3). Number of clusters were determined by using the optimal k elbow method within-clusters sum of squares (WCSS). Groups were classified into control (non-stressed), resilient (stressed), and susceptible (stressed) groups.

### Pavlovian discrimination paradigm and trial structure

In this Pavlovian paradigm, a highly palatable 30% sucrose solution (200 ms) served as the rewarding unconditioned stimulus (US), while a mildly punishment air puff to the subject’s face (∼10 psi, 100 ms) acted as the punishment US. Both the rewarding and punishment US were paired with a 5-second pure tone as the conditioned stimulus (CS), with the tone frequency set at 9 kHz for the rewarding CS and 2 kHz for the punishment CS. The reward trial started with the CS followed by a lick contingent reward US with a 2-second delay. After the CS ended, the US was vacuumed away from the spout. The punishment trials started with the CS followed by the punishment US with a 2-second delay. The reward and punishment catch trials both consisted of the respective CS with no US. The trials were separated by a 25-30 second inter-trial interval (ITI). Subjects were first head-fix trained in a closed box for 20 reward trials with no lick-contingency and no US delay. Each box was equipped with a replica of the acquisition setup, without the microscope. This consisted of a head-fix clamp fixed above the tube with the subject. A spout connected to a voltage recorder was fixed in front of the subject. The air puff spout and camera were fixed to opposite sides of the subject. Training sessions were ramped up to 60 trials over 3 sessions, after which lick contingency was turned on with a 2-second US delay for 2 sessions. Subsequently, Discrimination training sessions started, where 20% of trials changed to punishment trials. Before acquisition trials started subjects were trained under the 2-photon microscope for another 3 sessions. If subjects did not perform correctly anticipatory lick responses to > 50% of reward trials, learning was deemed unsuccessful. The acquisition sessions consisted of 8 punishment trials, 2 punishment catch trials (CS and no US), 36 reward trials without lick contingency, and 2 reward catch trials. These trials were pseudorandomized across the two blocks, with the requirements that the first 3 trials were reward trials, there were no consecutive sequences of 3 punishment trials, and the catch trials occurred in the last 15% of the trials. During each trial, facial footage, in vivo calcium imaging, and lick behavior was recorded.

### In vivo 2-photon calcium imaging

We used a two-photon microscope (Bruker Ultima Investigator, Bruker Nano) with a 20 × objective (0.45 NA, Olympus) and 920 nm excitation wavelength (Ti-Sapphire laser, Newport) for in vivo calcium imaging. Images were acquired using Prairieview (Bruker Nano) in resonant-galvo acquisition mode. Each field-of-view (FOV) (512 × 512 pixels covering 524 × 524 µm) was scanned at ∼29.8 Hz.

### Signal processing

Images from 2-photon calcium imaging were processed using Suite2P. We used Suite2P to correct motion artifacts, define regions of interest (ROIs) corresponding to individual neurons, and extract their GCaMP fluorescence (29). We selected only cellular ROIs by manual curation. Sessions and trials that contained motion artifacts and technical issues were taken out for further analysis. ROI match MATLAB software was used to identify cells that were successfully tracked across imaging sessions.

### Perfusion

Following the conclusion of recording experiments subjects were deeply anesthetized with an injection of sodium pentobarbital (200 mg/kg, intraperitoneal injection) and perfused transcardially with 20 mL of ice-cold lactated Ringer’s solution, followed by 20 mL ice-cold paraformaldehyde (4%; PFA) in phosphate-buffered saline (PBS). Brains were extracted and placed in 4% PFA for 24 h. The tissue was then equilibrated in a cryo-protectant solution (30% sucrose in PBS, w/v). Coronal slices measuring 60um were taken from the tissue using a sliding microtome (HM430; Thermo Fisher Scientific, Waltham, MA), and stored in PBS at 4 ^*°*^*C*.

### Epifluorescence imaging

Tissue slices were imaged using an epifluorescence microscope (Keyence BZ-X). Images were taken using a 2x objective lens. Following imaging, the images were evaluated to determine the location of viral expression as seen via GCaMP7f. Recording sites were located using GRIN lens lesion locations.

### Principal component analysis

Principal component analysis (PCA) was used to measure population firing rate dynamics in the mPFC (30). A local and global PCA was done on a matrix containing all z-scored normalized data (Reward CS tone, Punishment CS tone, Reward first lick, Reward US, Punishment US) for all animals such that we could compare neural trajectories across groups (control, resilient, and susceptible). For the local PCA, the matrix had neurons in rows, and in the columns had mean z-score response during-10 to 10 sec post CS event using 100 ms bins. The neural trajectories for each task-relevant event were created per group by multiplying the coefficients obtained in the PCA by the mean z-score response across trials per week. For each neural trajectory, the length was calculated as the sum of Euclidean distances between adjacent 100 ms bins. Also, neural trajectories distances were calculated as the Euclidean distance between the two trajectories bin-by-bin. For statistical comparison analysis, the neural trajectory metrics were calculated using the leave-one-out (LOO) method, leaving out all the neurons from a single animal per group, therefore the number of iterations is the number of mice in that group. Thus, in every iteration the same PCA coefficients per cell were used for neural trajectory analysis. For quantification of trajectory lengths and distance between trajectories the first 23 PCs were used to capture 59.51% of the variance. For all trajectory visualizations and trajectory quantifications, we matched the number of neurons for each group (control, resilient, and susceptible) for comparison analysis across weeks.

### Generalized linear model classifier

To test if anhedonia phenotype groups (control, resilient, and susceptible) could be decoded during reward and punishment trials from mPFC population activity, we used a generalized linear model (GLM) classifier (specifically, we applied multinomial logistic regression in MATLAB with default settings). To obtain anhedonia group mPFC population activity we used the coefficients obtained for each neuron in the local PCA and created a neural trajectory using the mean z-score responses for the Reward and Punishment trials (Reward first lick and Punishment US). We trained the GLM using the first 8 PCs per session per week (-10 to 10 seconds post CS event) as features. We performed a 10-fold cross-validation (CV), where the data was split into 10 subsets and in each iteration the training consisted of a different 90% subset of the data, then the testing was done with the remaining 10% of the data. For the 10-fold CV, we computed the area under the receiver operating characteristic curve (AUC score) for the test data. We used this model decode control versus resilient, control versus susceptible, and resilient versus susceptible. We then compared decoding performance (auROC scores) against shuffled data across weeks.

### Social analysis

To automatically detect social interaction behaviors, SLEAP (31) was used to estimate animal poses in behavior recordings. We recorded behavior videos using Noldus EthoVision XT and a Basler GenI Cam at 25 frames/second, set at a fixed distance above the three-chamber arena. A training data set was labeled using a 12-point skeleton to represent the mouse (nose, head, neck, left ear, right ear, left forepaw, right forepaw, left hindpaw, right hindpaw, trunk, tail base, tail tip), and was used to train a bottom-up model consisting of 2399 frames. To define interaction behavior with the social and nonsocial cups, we used a distance threshold of within 1.3x pixels to the radius of the cup and an angle threshold of 90 degrees between the subject’s nose, body, and the center of each cup to quantify time spent interacting across frames.

### Facial analysis

Video recordings of mouse facial expressions were collected on headfixed mice during discrimination sessions. We used SLEAP (31) version 1.2.9 (https://github.com/talmolab/sleap) to estimate the position of facial keypoints using a 13-point custom facial skeleton. This consisted of 4 points for eye (upper_eye, lower_eye, inner_eye, outer_eye,), 2 for whiskers (top_whisker_stem, bottom_whisker_stem), 4 for nose (nose_upper, nose_tip, nostril_left, nostril_right), and 3 for mouth area (mouth_upper, mouth_lower, chin). Our SLEAP model was trained on 11,154 manually labeled frames and consisted of a single-instance model with UNet backbone.

Analysis and visualizations were executed using MATLAB. We applied a smoothing filter to the SLEAP predictions using a Savitzky-Golay filter over a 5-frame window to minimize noise error associated with tracking. Using a custom built MATLAB toolbox called Facial Expression Feature Extractor (FEFE; https://github.com/Tyelab/FEFE), we extracted from the SLEAP pose estimates various facial features such as distances between keypoints, angles, velocities and accelerations of the nose and eye regions, and the areas of different facial regions as documented in Table 1. To reduce the bias of camera placement on our distance based features, we converted from pixels to cm by measuring the sucrose spout in each video and computing a pixel to cm conversion factor for that video.

**Table 1.**
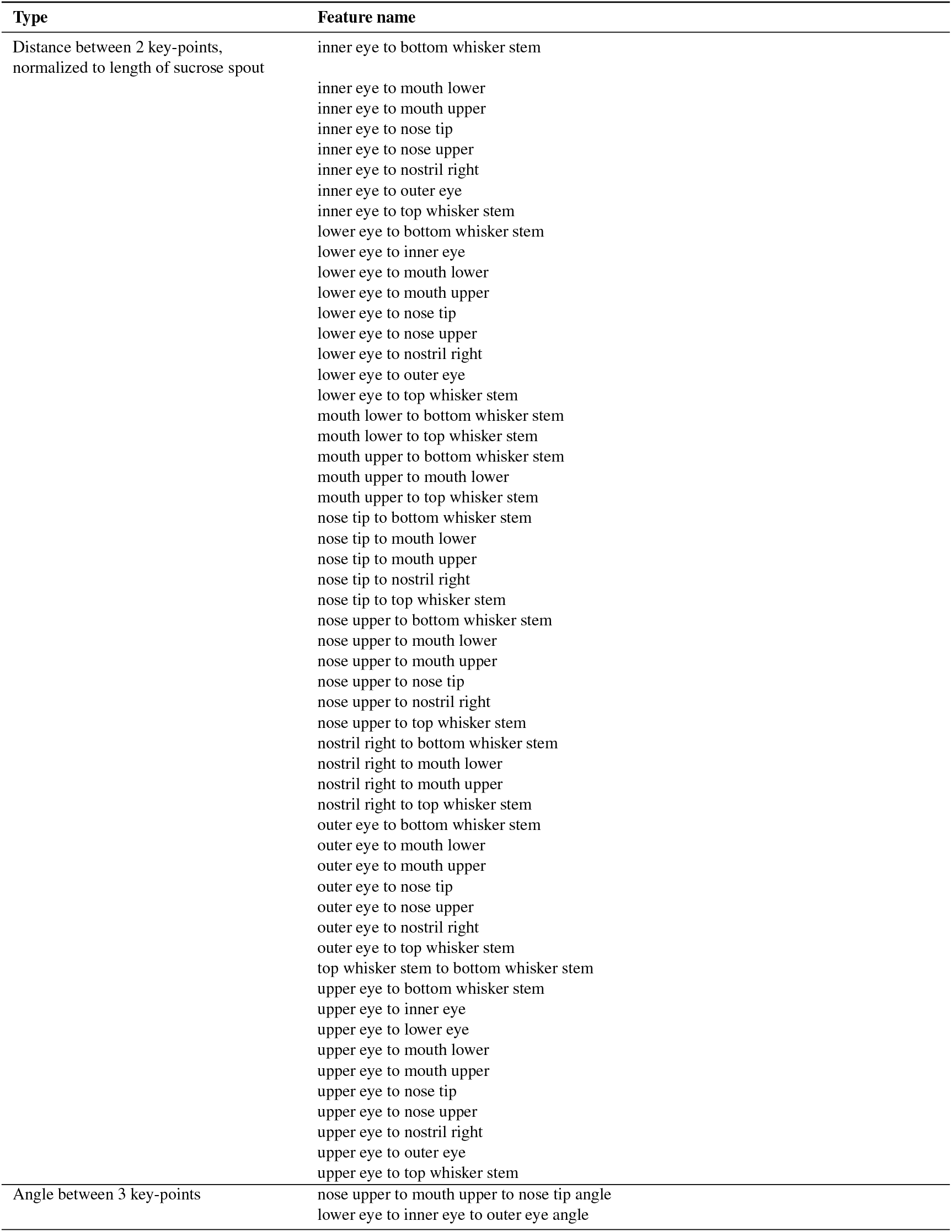
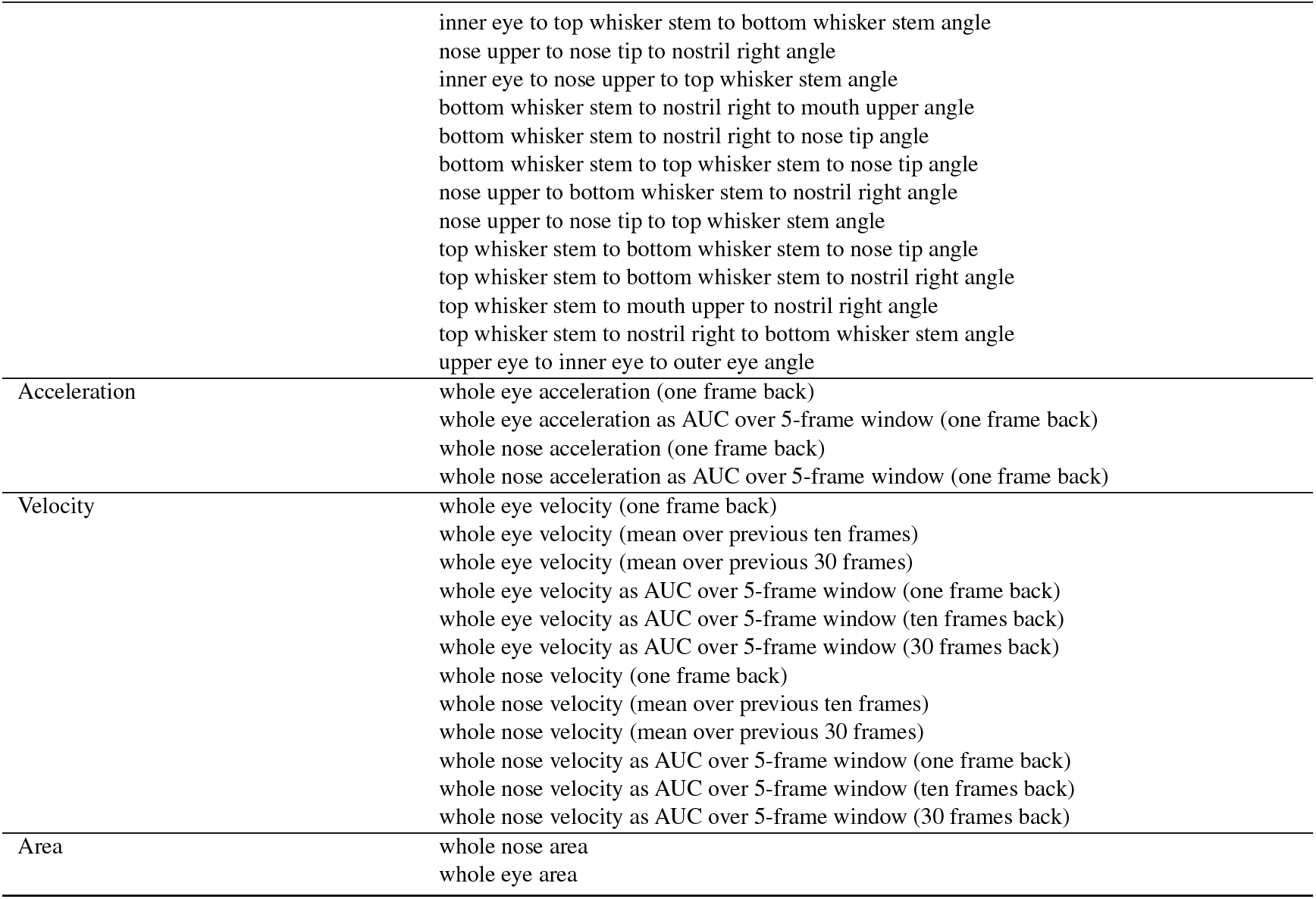
Table 1 Features computed and used for facial expression.

| Type | Feature name |
| --- | --- |
| Distance between 2 key-points,<br>normalized to length of sucrose spout | inner eye to bottom whisker stem |
|  | inner eye to mouth lower |
|  | inner eye to mouth upper |
|  | inner eye to nose tip |
|  | inner eye to nose upper |
|  | inner eye to nostril right |
|  | inner eye to outer eye |
|  | inner eye to top whisker stem |
|  | lower eye to bottom whisker stem |
|  | lower eye to inner eye |
|  | lower eye to mouth lower |
|  | lower eye to mouth upper |
|  | lower eye to nose tip |
|  | lower eye to nose upper |
|  | lower eye to nostril right |
|  | lower eye to outer eye |
|  | lower eye to top whisker stem |
|  | mouth lower to bottom whisker stem |
|  | mouth lower to top whisker stem |
|  | mouth upper to bottom whisker stem |
|  | mouth upper to mouth lower |
|  | mouth upper to top whisker stem |
|  | nose tip to bottom whisker stem |
|  | nose tip to mouth lower |
|  | nose tip to mouth upper |
|  | nose tip to nostril right |
|  | nose tip to top whisker stem |
|  | nose upper to bottom whisker stem |
|  | nose upper to mouth lower |
|  | nose upper to mouth upper |
|  | nose upper to nose tip |
|  | nose upper to nostril right |
|  | nose upper to top whisker stem |
|  | nostril right to bottom whisker stem |
|  | nostril right to mouth lower |
|  | nostril right to mouth upper |
|  | nostril right to top whisker stem |
|  | outer eye to bottom whisker stem |
|  | outer eye to mouth lower |
|  | outer eye to mouth upper |
|  | outer eye to nose tip |
|  | outer eye to nose upper |
|  | outer eye to nostril right |
|  | outer eye to top whisker stem |
|  | top whisker stem to bottom whisker stem |
|  | upper eye to bottom whisker stem |
|  | upper eye to inner eye |
|  | upper eye to lower eye |
|  | upper eye to mouth lower |
|  | upper eye to mouth upper |
|  | upper eye to nose tip |
|  | upper eye to nose upper |
|  | upper eye to nostril right |
|  | upper eye to outer eye |
|  | upper eye to top whisker stem |
| Angle between 3 key-points | nose upper to mouth upper to nose tip angle |
|  | lower eye to inner eye to outer eye angle |
|  | inner eye to top whisker stem to bottom whisker stem angle<br>nose upper to nose tip to nostril right angle<br>inner eye to nose upper to top whisker stem angle<br>bottom whisker stem to nostril right to mouth upper angle<br>bottom whisker stem to nostril right to nose tip angle<br>bottom whisker stem to top whisker stem to nose tip angle<br>nose upper to bottom whisker stem to nostril right angle<br>nose upper to nose tip to top whisker stem angle<br>top whisker stem to bottom whisker stem to nose tip angle<br>top whisker stem to bottom whisker stem to nostril right angle<br>top whisker stem to mouth upper to nostril right angle<br>top whisker stem to nostril right to bottom whisker stem angle<br>upper eye to inner eye to outer eye angle |
| Acceleration | whole eye acceleration (one frame back)<br>whole eye acceleration as AUC over 5-frame window (one frame back)<br>whole nose acceleration (one frame back)<br>whole nose acceleration as AUC over 5-frame window (one frame back) |
| Velocity | whole eye velocity (one frame back)<br>whole eye velocity (mean over previous ten frames)<br>whole eye velocity (mean over previous 30 frames)<br>whole eye velocity as AUC over 5-frame window (one frame back)<br>whole eye velocity as AUC over 5-frame window (ten frames back)<br>whole eye velocity as AUC over 5-frame window (30 frames back)<br>whole nose velocity (one frame back)<br>whole nose velocity (mean over previous ten frames)<br>whole nose velocity (mean over previous 30 frames)<br>whole nose velocity as AUC over 5-frame window (one frame back)<br>whole nose velocity as AUC over 5-frame window (ten frames back)<br>whole nose velocity as AUC over 5-frame window (30 frames back) |
| Area | whole nose area<br>whole eye area |

For the baseline session, we applied a local z-score to the set of 87 facial features, where the mean and standard deviation was computed for each trial using the 10 sec immediately preceeding that trial. We performed principal component analysis (PCA) on this set of facial features as follows: we concatenated the baseline data only into a matrix using normalized, trial-averaged facial features over an interval of-10 sec prior to CS-onset to +10 sec after CS-onset, yielding a matrix of 87 features by 29136 time points (607 time bins x 24 subjects x 2 trial types). Following PCA this was reduced to an 87×87 principal component coefficient matrix. Data from later weeks was projected into PC space using this baseline pca coefficient matrix using the first 10 PCs, which is associated with 91.0% of the variance explained. This process is notably different from principal component analysis of neural data because, unlike unlike neurons, the same facial features are able to be computed over all animals and all sessions.

For Stress sessions (weeks 1-6), we computed the mean and standard deviation of each facial feature using the entire baseline (Week 0) session and applied these values using a z-score to subsequent sessions. We perfomed PCA using only the baseline session data, as before, by concatenating the baseline data into a matrix using baseline-normalized, trial-averaged facial features over an interval of-10 sec prior to CS-onset to +10 sec after CS-onset, yielding a matrix of 87 features by 29136 time points (607 time bins x 24 subjects x 2 trial types).

For saline, ketamine, and post-ketamine sessions, we computed the mean and standard deviation of each facial feature using the post-stress session (Week 6) and applied these values using a z-score to subsequent sessions. We perfomed PCA using only the baseline session data, as before, by concatenating the post-stress data into a matrix using post-stress-normalized, trial-averaged facial features over an interval of-10 sec prior to CS-onset to +10 sec after CS-onset, yielding a matrix of 87 features by 29136 time points (607 time bins x 24 subjects x 2 trial types). Data from saline and ketamine administration sessions, and from post-ketamine sessions, were normalized as described, then projected into the post-stress PCA space.

To display PCA, we averaged across trajectories for each subject within each phenotype. To compute trajectory lengths, we computed the Euclidean norm of each subject’s trajectory, then took the mean across subjects. For distance between trajectories, we took the Euclidean norm of the pointwise differences of sucrose and air puff trajectories for each time step for each session; from this we also computed average distance by phenotype.

For facial decoding, we projected the data into PCA space, then applied a multinomial logistic regression model. We used a 10-fold cross validation and compared the results to a control model where the phenotype labels were shuffled in random order. The area under the curve (AUC) metric was smoothed by applying a Gaussian moving average in a window using the previous 20 sec.

### Statistical methods

The thresholds for significance were placed at *p<0.05, **p<0.01, ***p<0.001, and ****p<0.0001 unless stated otherwise. All data are shown as mean and SEM. Wilcoxon signed rank-sum test, Pearson correlation, one-way ANOVA, Repeated-measure ANOVA, and mixed-effects model followed by a Tukey’s post-hoc test, two-way ANOVA followed by Dunnet’s multiple comparison test, and Forsythe-Brown test and Welch’s ANOVA were performed using GraphPad Prism 6, GraphPad Prism 10, or MATLAB. Data was checked for adherence to normality. Any outliers were removed using Grubb’s Test. The p values were corrected for multiple comparisons. Ward’s linkage hierarchical clustering utilizing Euclidean distance was performed using MATLAB.

**Extended Data Figure 1.**
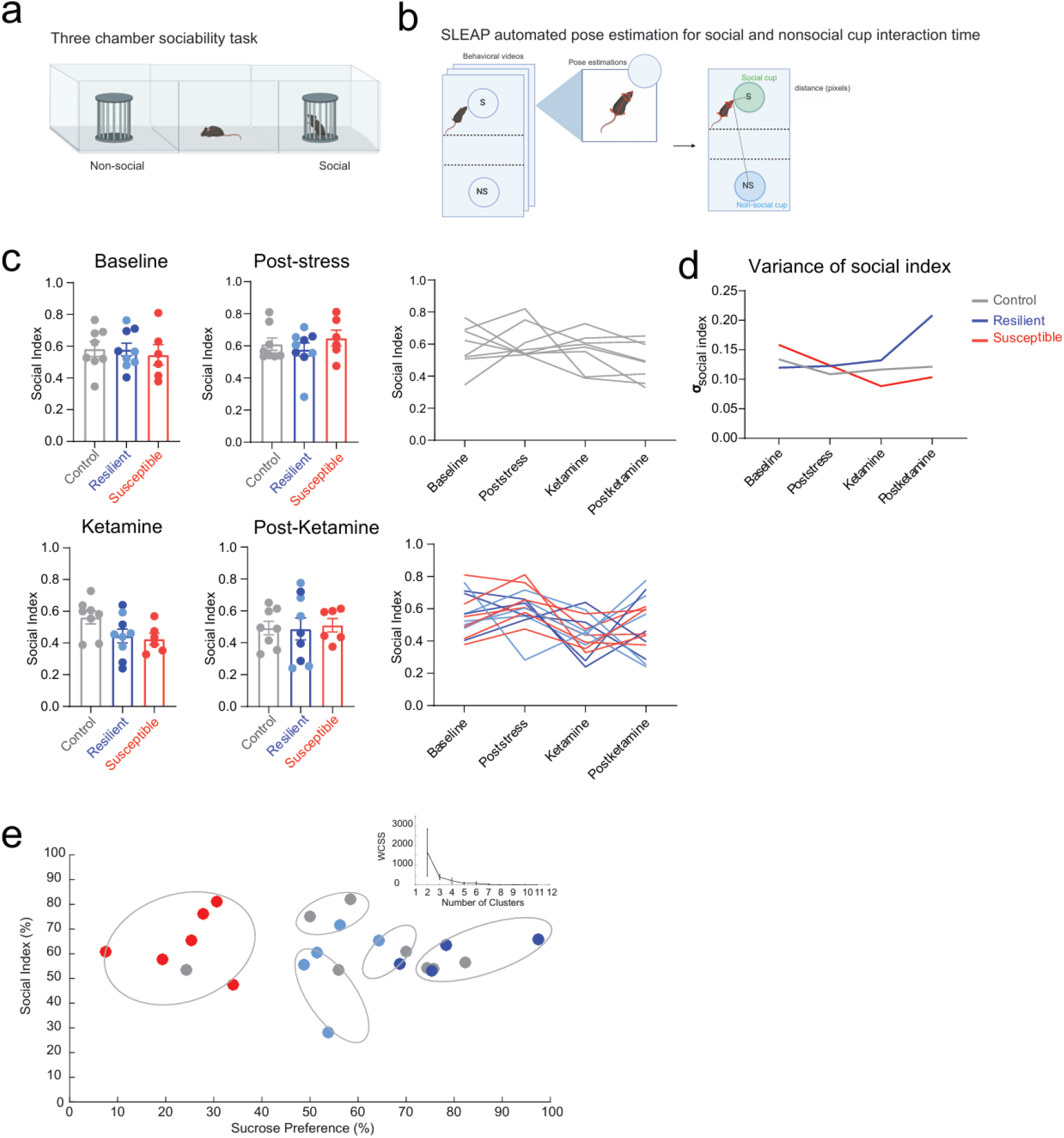
Ketamine treatment after chronic mild stress decreases variance in social index in susceptible mice. **a**. Schematic of three-chamber sociability task assessing social preference. **b**. Workflow for SLEAP automated pose tracking, used to precisely quantify interaction time based on the subject’s distance in pixels and angle to both the social and non-social cups. **c**. No difference in social interaction across groups at Baseline, Post-stress, ketamine, and Post-Ketamine time points. Social index, calculated as a ratio of time spent interacting with the social cup over combined social cup and non-social cup interaction times, measured at Baseline (one-way ANOVA, Tukey’s post hoc, interaction effect: F _(2, 20)_ = 0.1539, p=0.8583), Post-stress (one-way ANOVA, Tukey’s post hoc, interaction effect: F _(2, 20)_=0.09649, p=0.5403), Ketamine (one-way ANOVA, Tukey’s post hoc, interaction effect: F _(2, 20)_=0.2762, p=0.0726), and one week after Ketamine treatment (one-way ANOVA, Tukey’s post hoc, interaction effect: F _(2, 20)_ =3.173, p=0.9614) time points. Error bars represent mean +/-SEM. **d**. Standard deviation plot of social index across Baseline (control: n=8, SD= 0.1338; resilient: n=9, SD=0.1196; susceptible: n=5, SD=0.1585), Post-stress (control: n=8, SD=0.1087; resilient: n=9, SD=0.1255; susceptible: n=5, SD=0.1234), Ketamine (control: n=8, SD=0.1166; resilient: n=9, SD=0.1322; susceptible: n=5, SD=0.08842), and after Ketamine (control: n=8, SD=0.1213; resilient: n=9, SD=0.2082; susceptible: n=5, SD=0.1036) time points. **e**. k-means clustering (k=5) of social index and sucrose preference scores. The optimal k elbow method using the within-cluster-sum-of-square (WCSS) was applied to determine the appropriate number of clusters derived from social index and sucrose preference scores of mice Post-stress time point.

**Extended Data Figure 2.**
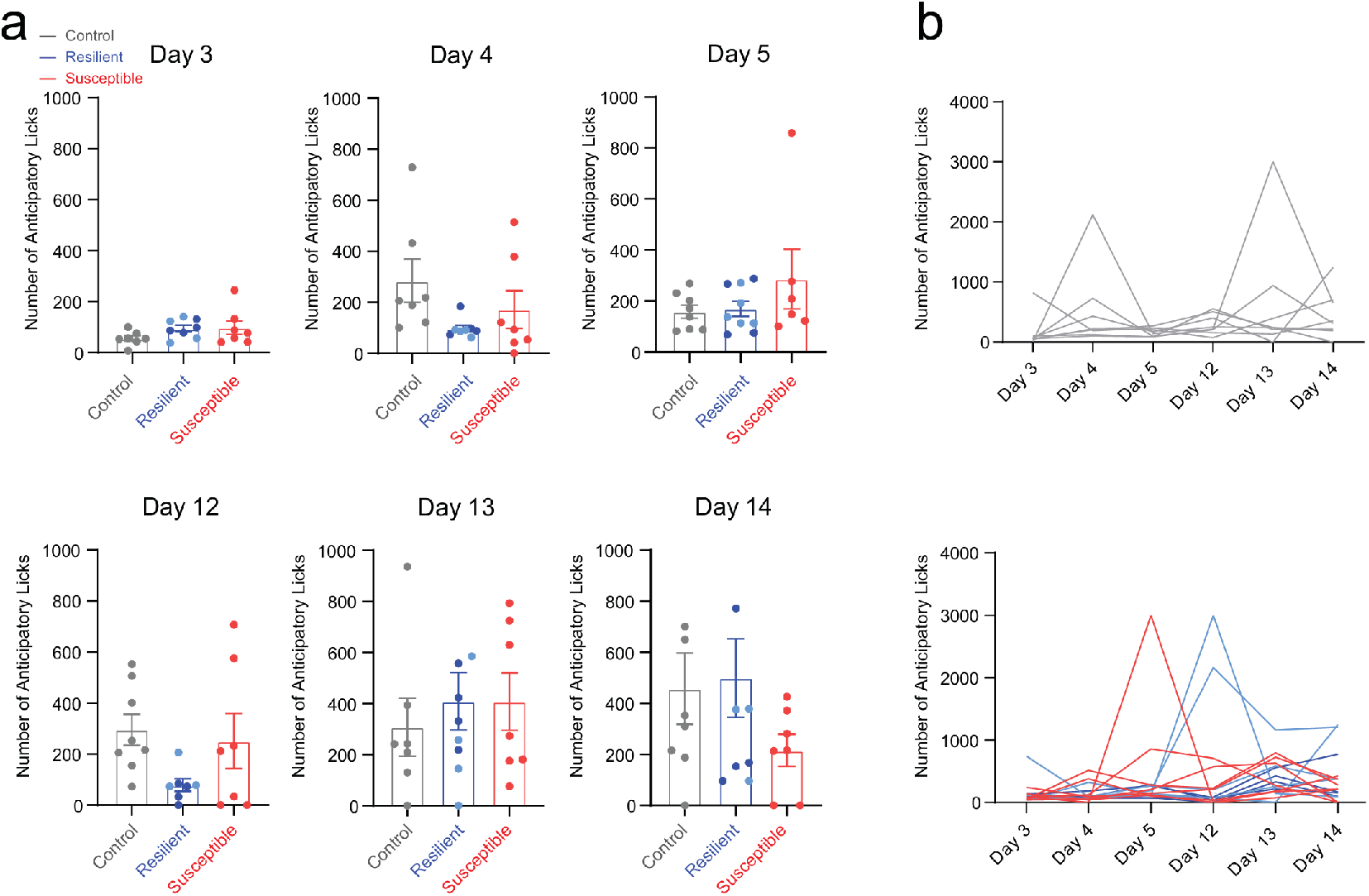
Susceptible mice show no differences in anticipatory licking during head-fixed training task prior to stress. **a**. During head-fixed training, total number of anticipatory licks (CS-onset at 0 sec to US-delivery at 2 sec) measured at multiple time points. No significant differences across control, resilient, and susceptible groups: day 3 (one-way ANOVA, Tukey’s post hoc, interaction effect: F _(2, 19)_ =1.644, p=0.2196), day 4 (one-way ANOVA, Tukey’s post hoc, interaction effect: F _(2, 19)_ =2.353, p=0.1221), day 5 (one-way ANOVA, Tukey’s post hoc, interaction effect: F _(2, 20)_ = 1.295, p=0.2958), day 12 (one-way ANOVA, Tukey’s post hoc, interaction effect: F _(2, 19)_ =2.520, p=0.1070), day 13 (one-way ANOVA, Tukey’s post hoc, interaction effect: F _(2, 20)_=0.2470, p=0.7835), and day 14 (one-way ANOVA, Tukey’s post hoc, interaction effect: F _(2, 21)_ = 1.249, p=0.3073) of headfixed training. Error bars represent mean +/-SEM. **b**. Longitudinal description showing non-stressed control mice (top panel: gray) and stressed (bottom panel: resilient and susceptible) mice during headfixed training.

**Extended Data Figure 3.**
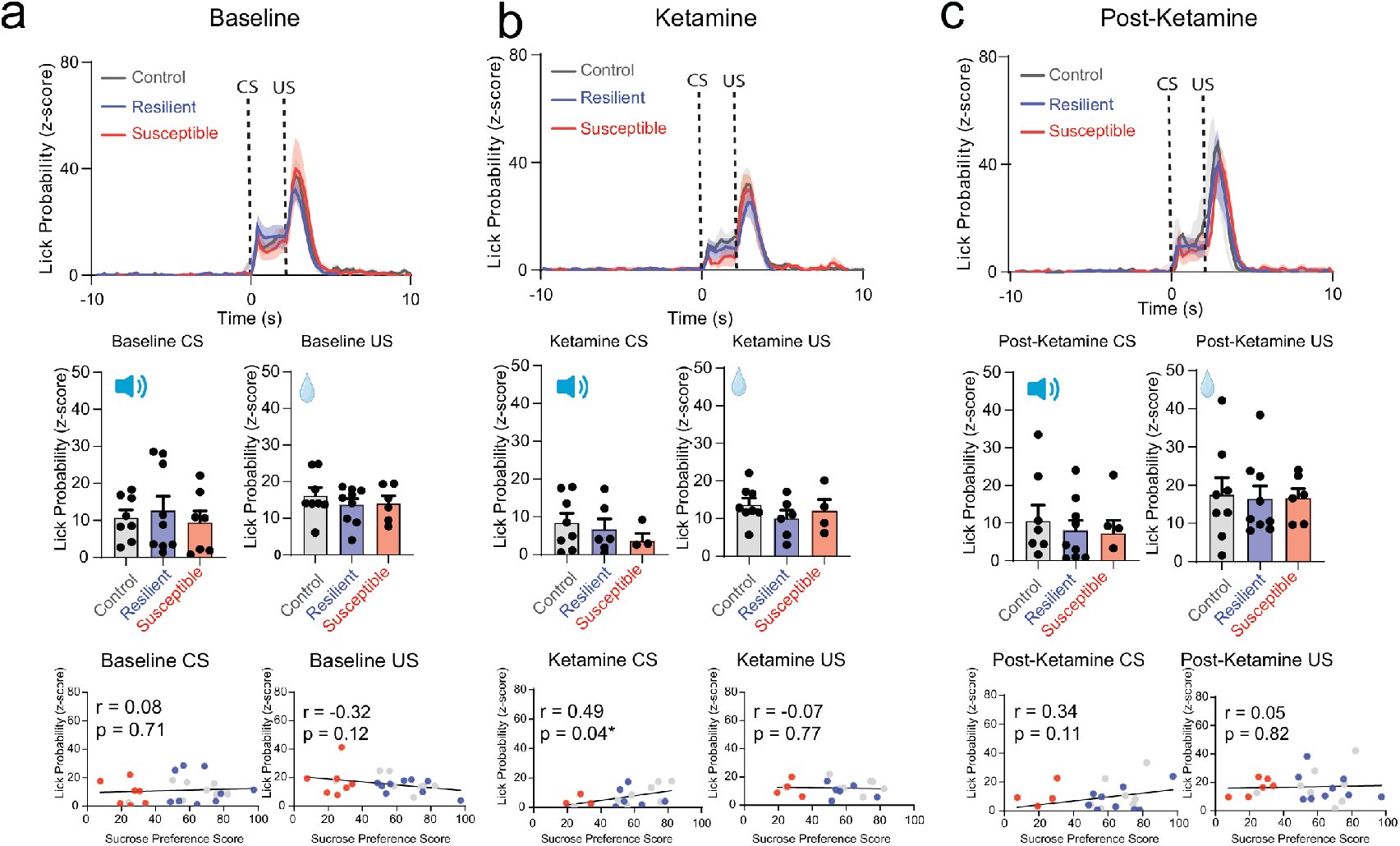
No difference in lick probability within susceptible group at Baseline, Ketamine, and Post-Ketamine time points. **a**. Visualizing lick probability relative to cue onset of CS (0 - 2 seconds) and sucrose delivery of US (2 - 5 seconds) in control, resilient and susceptible groups during Baseline (top panel). No significant differences in lick probability across groups: Lick Probability (One-way ANOVA, Baseline CS, F_(2, 20)_= 0.5011, p= 0.6133; Baseline US, F_(2, 20)_= 0.4939, p= 0.6175) (middle panel).). No correlation in lick probability and sucrose preference at Baseline. Pearson’s correlation of lick probability and sucrose preference test Baseline CS r= 0.08, p= 0.71, Baseline US r=-0.32, p= 0.12. (bottom panel). **b**. No significant differences in lick probability across groups: Lick probability relative to cue onset of CS and sucrose delivery of US in control, resilient and susceptible groups during Ketamine time point (top panel). Lick Probability (One-way ANOVA, Ketamine CS, F_(2, 15)_= 0.8240, p= 0.4576; Ketamine US, F_(2, 20)_= 0.2545, p=0.7778) (middle panel). Significant correlation in lick probability and sucrose preference at Ketamine time point during CS, but not US. Pearson’s correlation of lick probability and sucrose preference test Ketamine CS r= 0.49, p= 0.039* Ketamine US r=-0.07, p= 0.77 (bottom panel). **c**. No significant differences in lick probability across groups: Lick probability relative to cue onset of CS and sucrose delivery of US in control, resilient and susceptible groups during post-Ketamine timepoint (top panel). Lick Probability (One-way ANOVA, post-Ketamine CS, F_(2, 20)_= 0.0239, p=0.9764. (middle panel). No correlation in lick probability and sucrose preference at post-Ketamine. Pearson’s correlation of lick probability and sucrose preference test post-Ketamine CS r= 0.34, p= 0.11, post-Ketamine US r= 0.05, p= 0.82 (bottom panel).

**Extended Data Figure 4.**
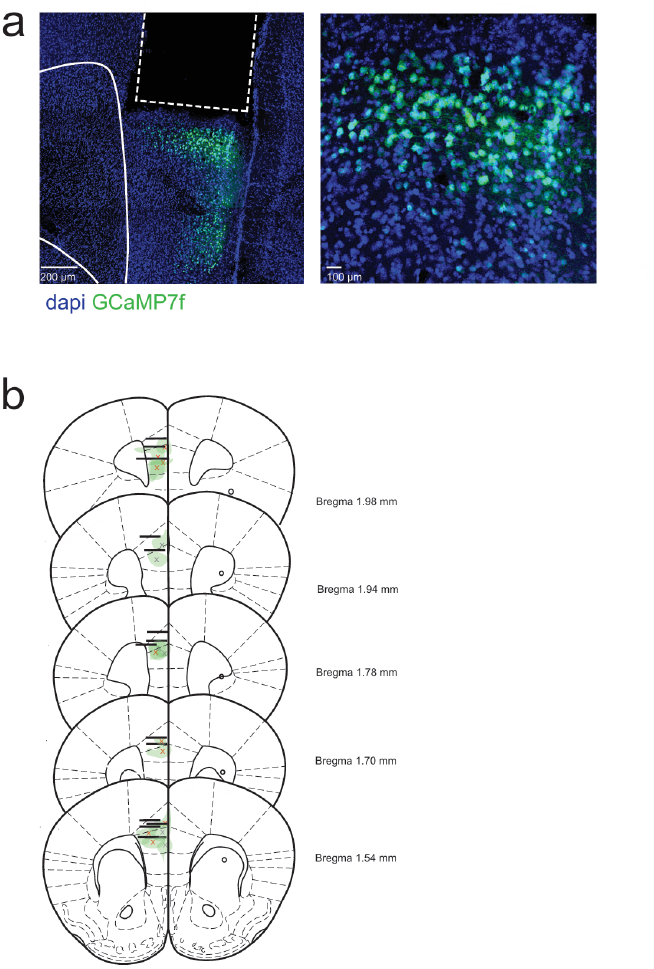
Histological validation of injection sites and implants. **a**. Representative images of GRIN lens implant and GCaMP7f expression in the PFC **b**. GRIN lens implant locations and GCaMP7f injection sites in the mPFC for in vivo 2-photon calcium recording (Bregma 1.54 to 1.98 mm). x indicates viral injection site.

**Extended Data Figure 5.**
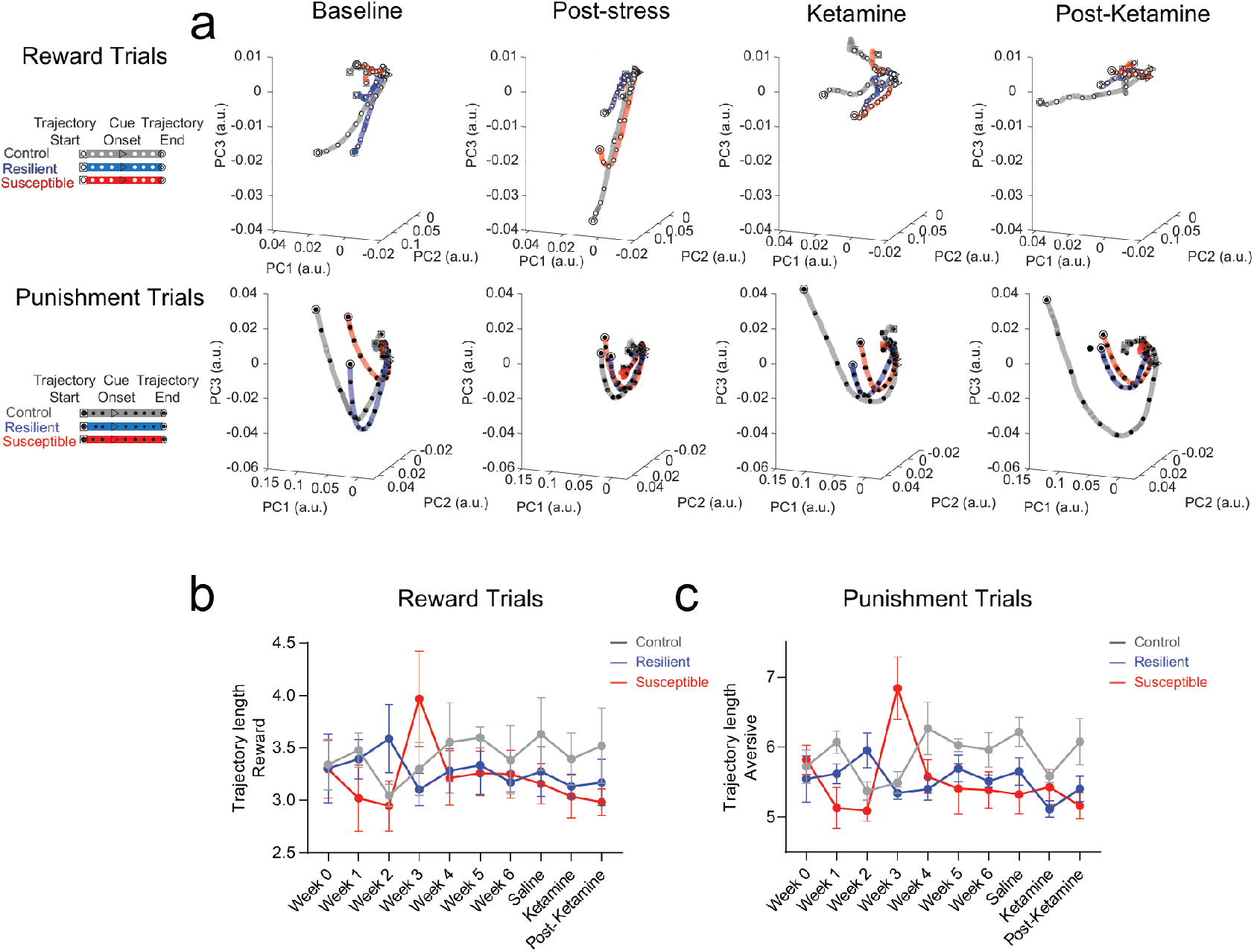
Visualizing neural population activity as neural trajectories using local Z-score revealed no differences across weeks. **a**. Neural trajectory lengths (post-event, 0-10 sec) in control, resilient, and susceptible groups during reward trials and punishment trials (using principal components that captured 90% of variance) across weeks. **b**. Reward (Left panel): Mixed ANOVA: subjects, F_(1.928, 109.9)_=8.184, p=0.0006, weeks, F_(9,114)_=1.638, p=0.1127, interaction, F_(18,114)_=4.126. c. Punishment (Right panel): subjects, F_(1.984, 113.1)_=8.475, p=0.0004, weeks, F_(9,114)_=1.154, p=0.3313, p<0.0001, interaction, F_(18,114)_=3.140, p=0.0001.

**Extended Data Figure 6.**
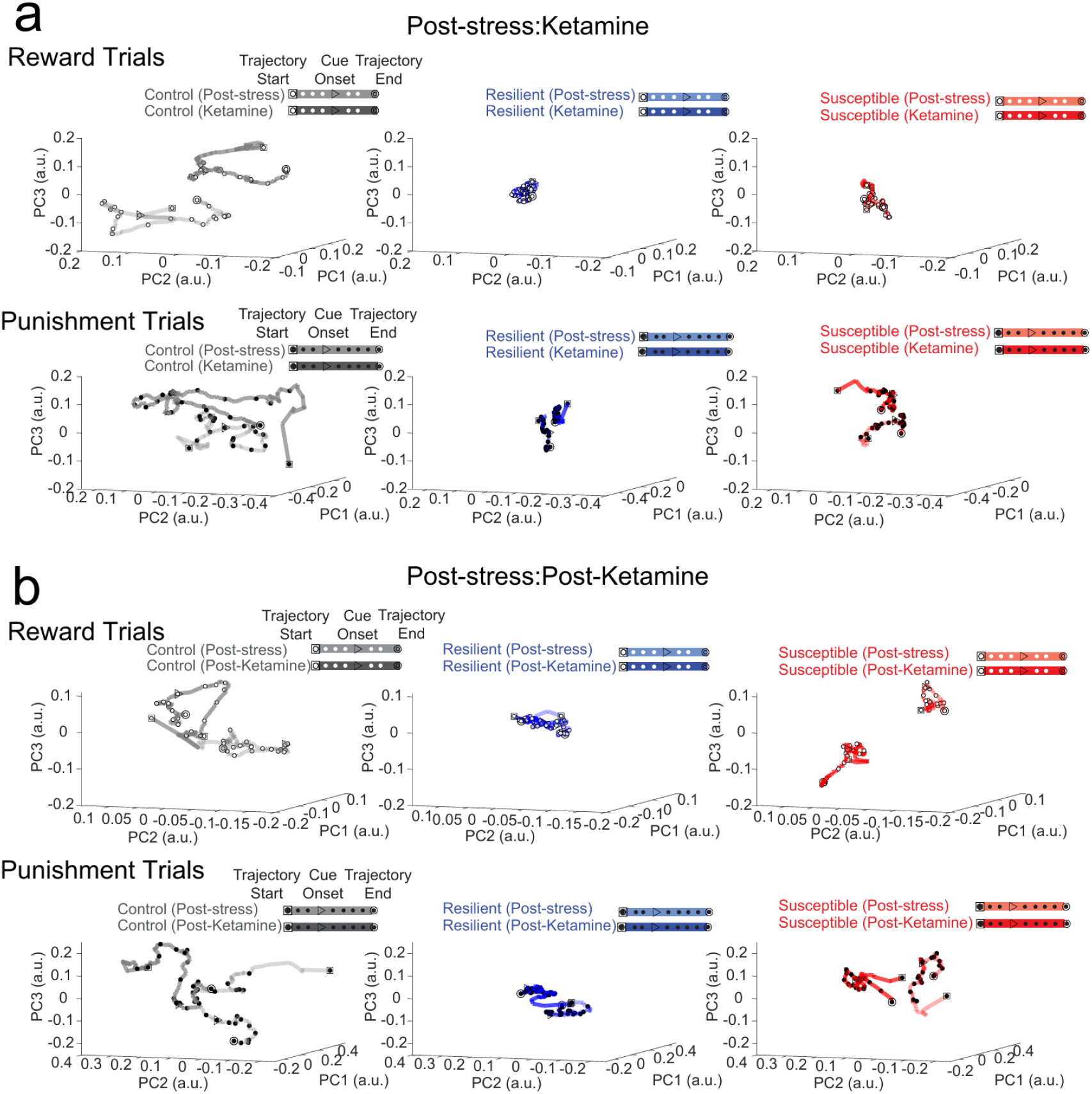
Neural trajectories of longitudinally imaged ensembles during Post-stress to Ketamine time points, and Post-stress to Post-Ketamine time points. **a**. Using neural trajectories of mPFC neural populations plotted with a super global Z-score (Z-score normalized across multiple sessions), ROI-matched populations between sessions during reward (Top) and punishment trials (Bottom) at Post-stress and Ketamine time points. **b**. ROI matched neural trajectories of mPFC neural populations during reward (Top) and punishment trials (Bottom) at Post-stress and Post-Ketamine time points.

**Extended Data Figure 7.**
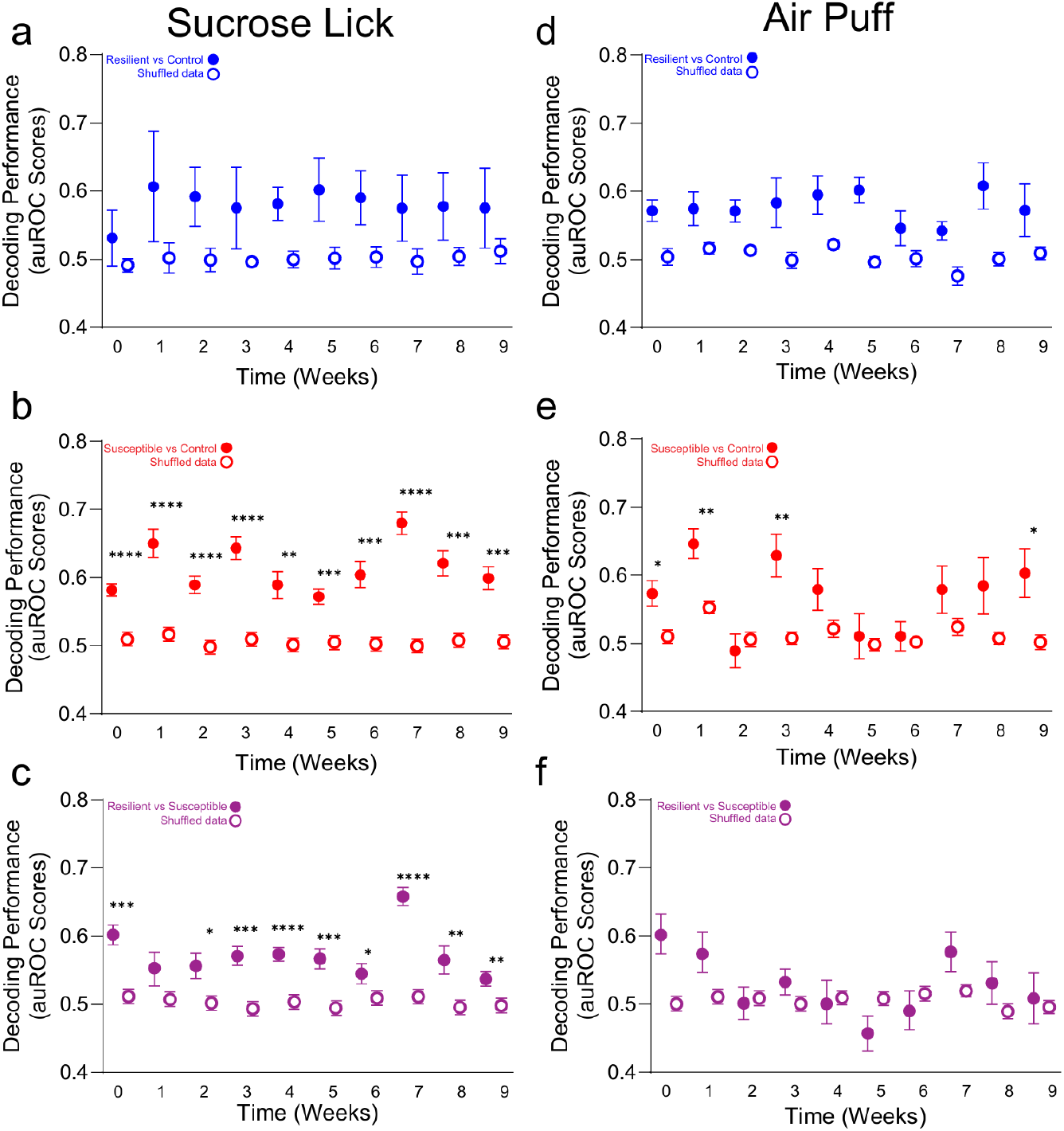
mPFC population activity decodes stress phenotypes. **a**. Significant decoding performance of resilient vs. control groups compared to shuffled data. Decoding accuracy in response to sucrose lick in resilient vs. control groups across weeks. Two-way Repeated Measures ANOVA, event F_(1,18)_=143.5, p<0.0001, weeks F_(3.899,70.17)_=2.145, p=0.0858, interaction F_(9,162)_=1.309, p=0.2361. **b**. Significant decoding performance of susceptible vs. control groups compared to shuffled data within individual weeks. Decoding accuracy in response to sucrose lick in *susceptible vs control* groups across weeks. Two-way Repeated Measures ANOVA, event F_(1,18)_=197.2, p<0.0001, weeks F_(3.788,68.18)_=4.813, p=0.0021, interaction F_(9,162)_=4.230, p<0.0001. Tukey post-hoc, Weeks 0-9: p<0.0001, p<0.0001, p<0.0001, p<0.0001, p=0.0014, p=0.0001, p=0.0004, p<0.0001, p=0.0001, p=0.0003. **c**. Significant decoding performance of resilient vs. susceptible groups compared to shuffled data within individual weeks. Decoding accuracy in response to sucrose lick in resilient vs. susceptible groups across weeks. Two-way Repeated Measures ANOVA, event F_(1,18)_=234.8, p<0.0001, weeks F_(4.550,81.91)_=5.171, p=0.0005, interaction F_(9,162)_=3.633, p=0.0004. Tukey post-hoc, Weeks 0-9: p=0.0001, p=0.1186, p=0.0171, p=0.0003, p<0.0001, p=0.0007, p=0.0384, p<0.0001, p=0.0081, p=0.0055. **d**. No significant difference in decoding performance of resilient vs. control groups compared to shuffled data across weeks. Decoding accuracy in response to air puff in resilient vs control groups across weeks. Two-way Repeated Measures ANOVA, event F_(1,18)_=72.28, p<0.0001, weeks F_(4.292,77.26)_=1.041, p=0.3943, interaction F_(9,162)_=0.5241, p=0.8556. **e**. Significant decoding performance of susceptible vs. control groups compared to shuffled data within individual weeks, but not week 2, and weeks 4-8. Decoding accuracy in response to air puff in susceptible vs. control groups across weeks. Two-way ANOVA, event F_(1,18)_=51.47, p<0.0001, weeks F_(5.353,96.35)_=3.086, p=0.0028, interaction F_(9,162)_=1.883, p=0.0579. Tukey post-hoc, Weeks 0-9: p=0.0105, p=0.0017, p=0.5491, p=0.0036, p=0.1050, p=0.7347, p=0.7196, p=0.1682. **f**. No significant difference in decoding performance of resilient vs. susceptible groups across weeks. Decoding accuracy in response to air puff in resilient vs. susceptible groups across weeks. Two-way Repeated Measures ANOVA, event F_(1,18)_=7.780, p=0.0121, weeks F_(4.555,81.99)_=2.203, p=0.0676, interaction F_(9,162)_=2.225, p=0.0229. All post-hoc comparisons are Tukey t-tests, *p<0.05, **p<0.01, ***p<0.001, ****p<0.0001 All 2-way ANOVAs were for event (event vs. shuffle) and weeks (0-9).

**Extended Data Figure 8:**
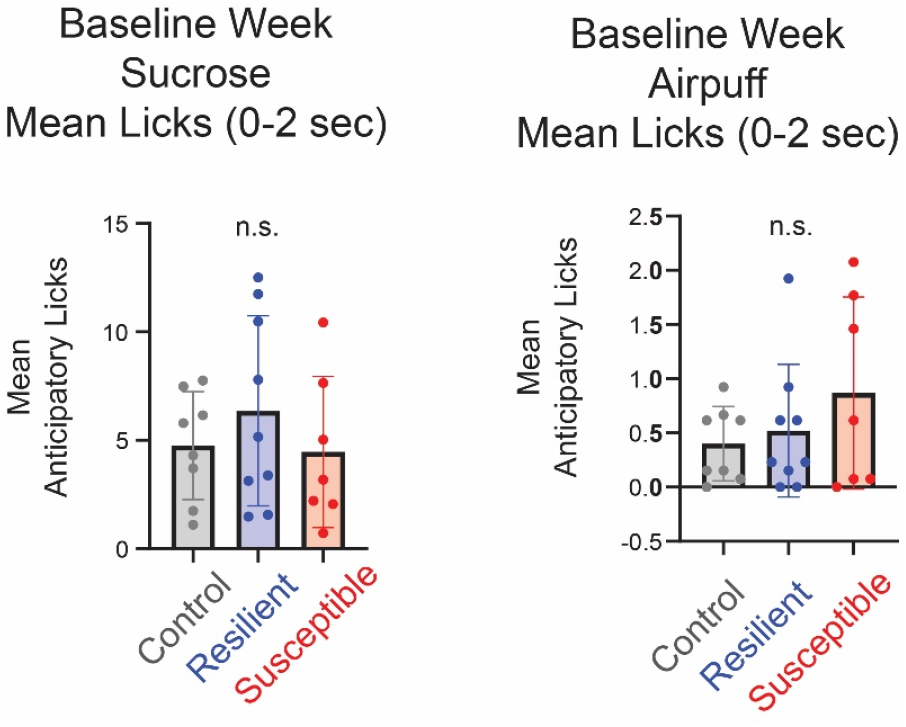
Mean Number of Anticipatory Licks during 0-2s in Week 0. There is no difference in mean number of anticipatory licks during the interval from CS-onset to just before US-delivery during Baseline week for any phenotype during Sucrose trials (left) or Air puff trials (right). (Sucrose, Ordinary one-way ANOVA, F_(2,21)_=0.6754, p=0.5201; Air puff, Kruskal-Wallis test, H = 0.3475, p=0.8405)

**Extended Data Figure 9:**
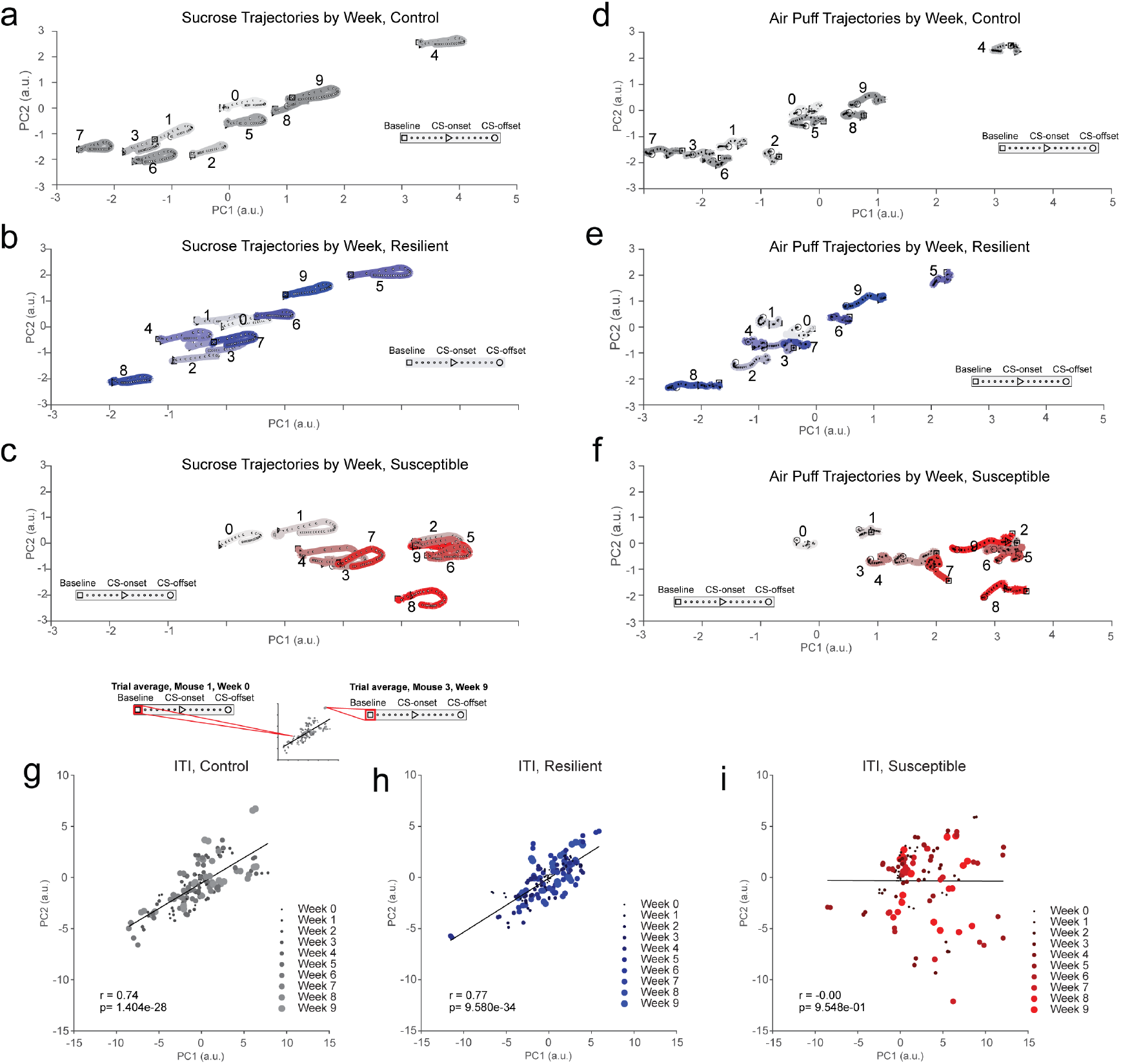
Significant correlation found between first two PCs in Resilient and Control Mice but not Susceptible Mice (a-f): Trajectories for facial feature dynamics for all weeks of each phenotype are shown from-10 s to +10 s interval around CS-reward onset at 0. Facial features were normalized to baseline session (week 0). We performed PCA on baseline session (week 0), then used the first 10 PCs (capturing 91% of variance explained) to project the remaining weeks into the same PC space as the baseline session. Each trajectory represents an average across trajectories from each subject in phenotype for the week shown. **a**. Trajectory plots showing (-10 s, +10 s) interval around CS-reward onset, normalized to baseline (Week 0), averaged across subjects in control group for all weeks. The baseline trajectory is located near (0, 0) in PC space and the other weeks spread out along the diagonal in an approximately normal distribution along the diagonal and around the baseline trajectory. **b**.Trajectory plots showing (-10 s, +10 s) interval around CS-reward onset, normalized to baseline (Week 0), averaged across subjects in resilient group for all weeks. The baseline trajectory is located near (0, 0) in PC space and the other weeks spread out along the diagonal in an approximately normal distribution along the diagonal and around the baseline trajectory. **c**. Trajectory plots showing (-10 s, +10 s) interval around CS-reward onset, normalized to baseline (Week 0), averaged across subjects in susceptible group for all weeks. The baseline trajectory is located near (0, 0) in PC space and the other weeks show a drift to the right along PC1 and slightly down in PC2, unlike the pattern shown in **a, b. d**. Trajectory plots showing (-10 s, +10 s) interval around CS-punishment onset, normalized to baseline (Week 0), averaged across subjects in control group for all weeks. The baseline trajectory is located near (0, 0) in PC space and the other weeks spread out in an apparently normal distribution along the diagonal and around the baseline trajectory. **e**. Trajectory plots showing (-10 s, +10 s) interval around CS-punishment onset, normalized to baseline (Week 0), averaged across subjects in resilient group for all weeks. The baseline trajectory is located near (0, 0) in PC space and the other weeks spread out along the diagonal in an approximately normal distribution along the diagonal and around the baseline trajectory. **f**. Trajectory plots showing (-10 s, +10 s) interval around CS-punishment onset, normalized to baseline (Week 0), averaged across subjects in susceptible group for all weeks. The baseline trajectory is located near (0, 0) in PC space and the other weeks show a drift to the right along PC1 and slightly down in PC2, unlike the pattern shown in **d, e. (g-i)**. To quantify the linear relationship between PC1 and PC2 observed in resilient and control groups, we extracted facial features z-scored to global baseline at a point during the intertrial interval (10 sec before CS-onset) for reward and punishment trials. Each point in the plot represents the projection of a subject’s trial averaged facial features for a given week onto the principal component space. **g**. There is significant correlation between PC1 and PC2 in the control group for all subjects across the 10 weeks (Pearson’s correlation coefficient R=0.74, p<0.0000****). **h**. Facial features from resilient mice show a strong correlation between PC1 and PC2 (Pearson’s correlation coefficient R=0.77, p<0.0000****). **i**. Susceptible animals, unlike control and resilient animals, do not exhibit detectable correlation between PC1 and PC2 in their facial features across the 10 weeks (Pearson’s correlation coefficient r=0.00, p=0.9548).

**Extended Data Figure 10:**
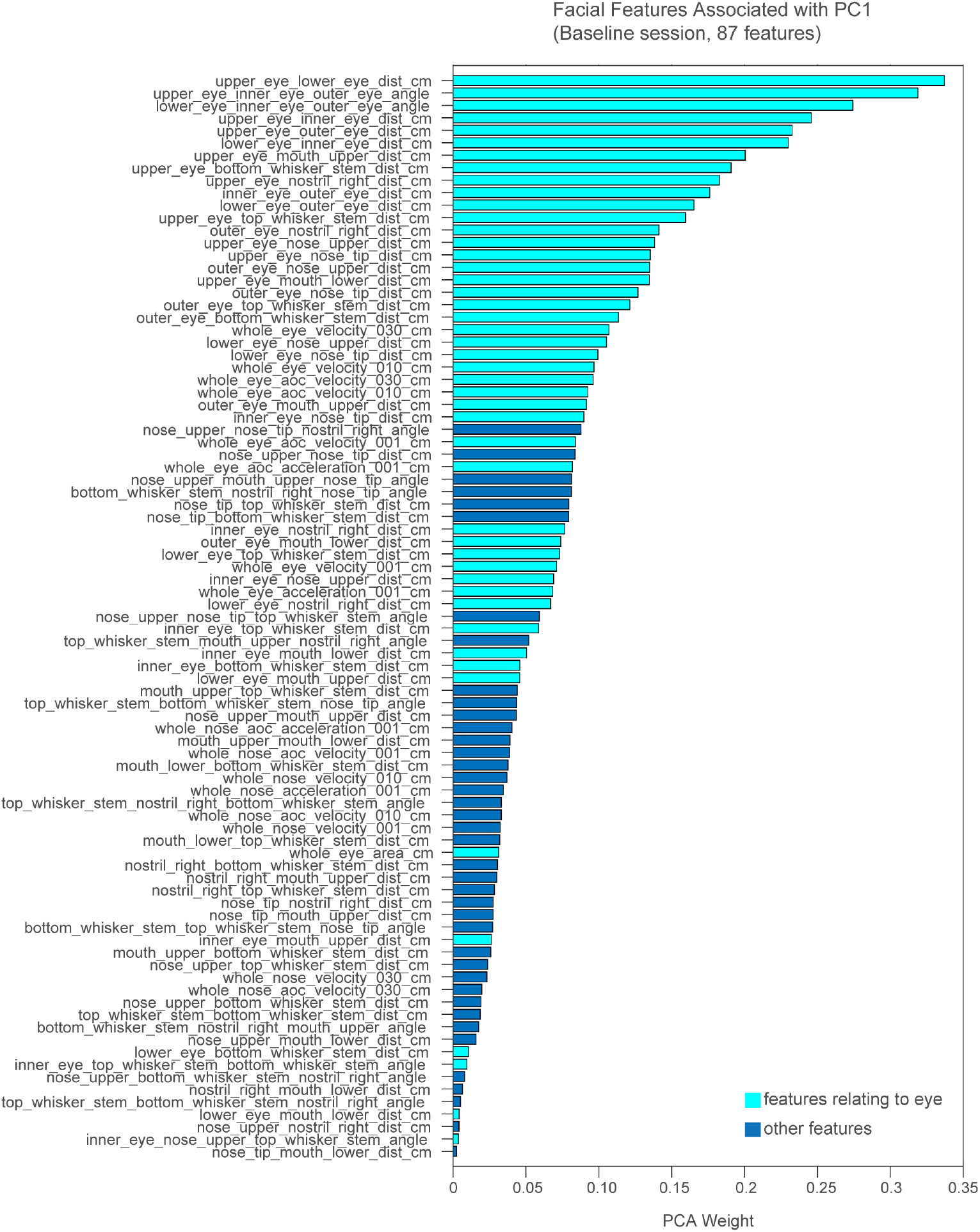
Eye-related Features are prominent in PC1. To explore the interpretation of the first principal component, we sorted the absolute value of the PCA weights associated with each facial feature and found that the highest weighted feature was the distance between the upper and lower eye, effectively describing eye-blinks. Cyan bars represent all features that include at least one eye key point in the feature name (for distances and angles) or other eye-related feature (area, velocity, or acceleration). Most of the highest weights in PC1 are associated with eye-related features.

**Extended Data Figure 11:**
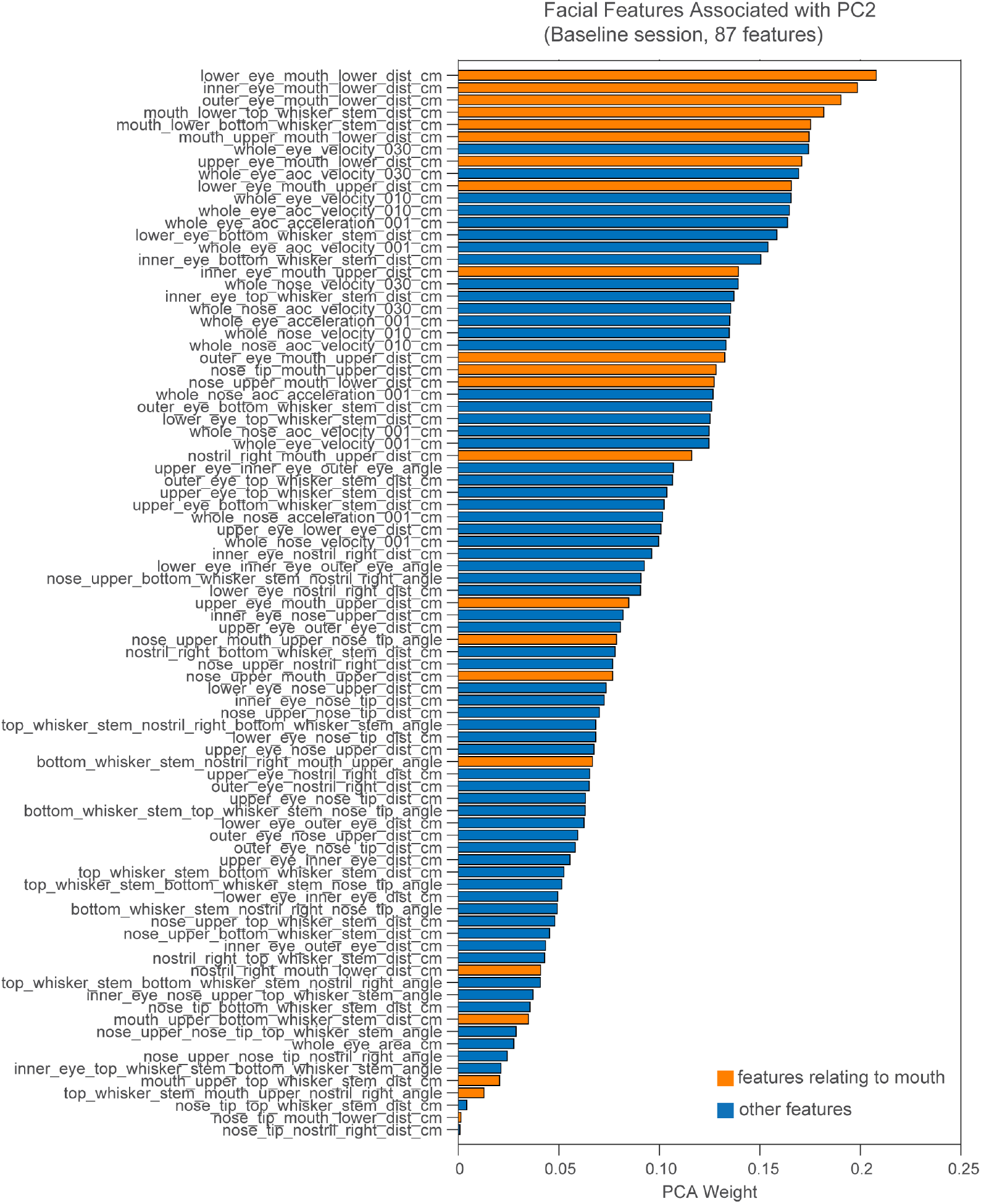
Mouth-related Features are prominent in PC2. To understand how to interpret the second principal component, we sorted the absolute value of the PCA weights associated with each facial feature and found that the first three highest features related to the distance between the eye and the lower mouth. In a head-fixed animal, the lower eye, inner eye, and outer eye points are stable when the eye is not blinking. We color-coded orange all features that include at least one mouth key point in the feature name and found that the top weighted features represent various distances involving the mouth. We therefore interpret PC2 as describing mouth movements, specifically the opening and closing of the mouth.

**Extended Data Figure 12.**
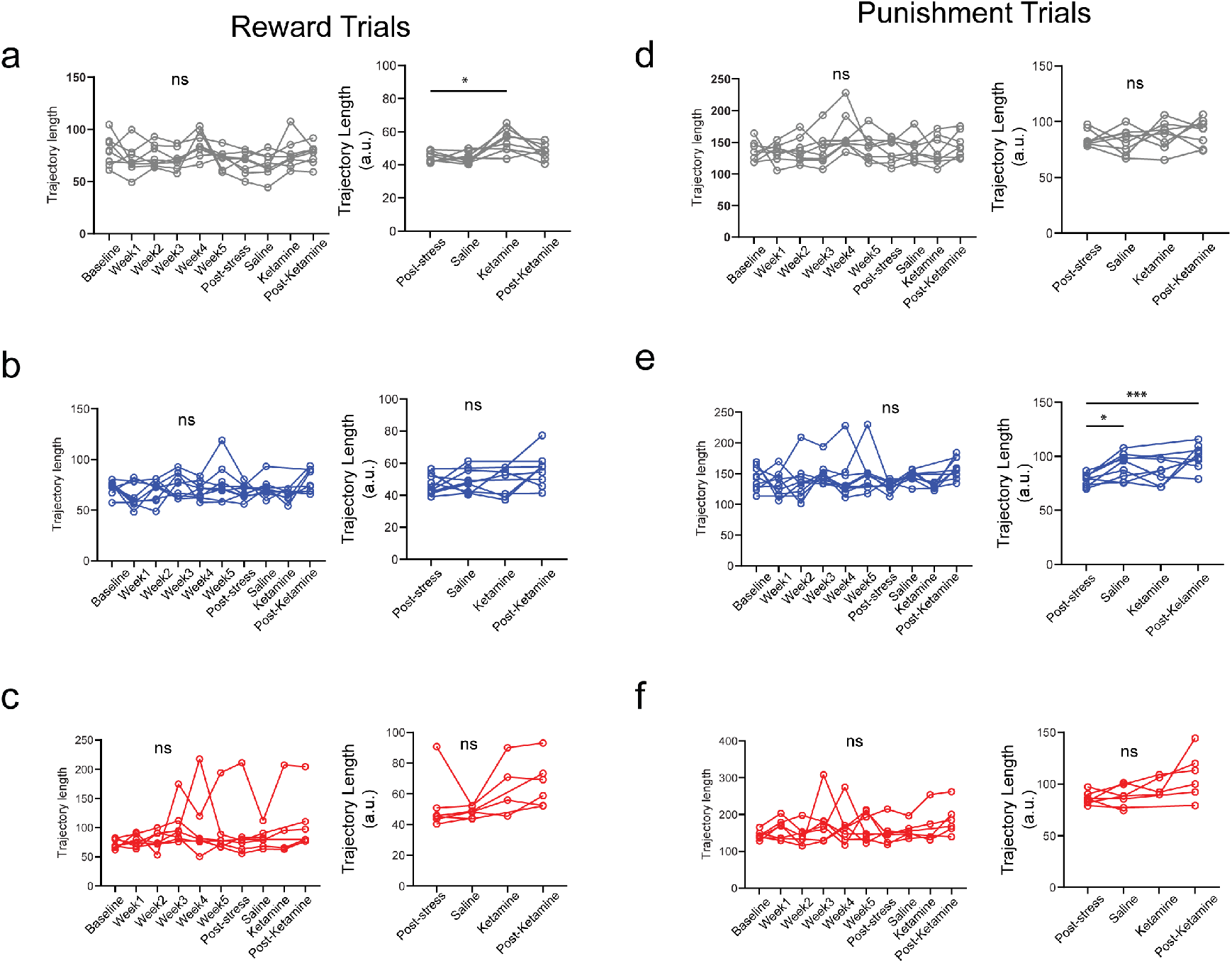
Control mice display increase in facial feature dynamics in response to reward during Ketamine week while resilient and susceptible mice do not. **a**. Control animals trajectory lengths post-event (0-10 sec at onset of CS) during reward trials, normalized to week 0 (left) are not significantly different across weeks but when normalized to post-stress Week 6 (right), the facial dynamics show an increase in ketamine week compared to post-stress (RM one-way ANOVA Week 0 normalization F_(3.499, 24.49)_ = 2.706, p=0.0600,; Mixed ANOVA week F _(1.981, 13.87)_ = 7.756, p=0.0056*). **b**. Resilient animals trajectory lengths post-event (0-10 sec at onset of CS) during reward trials, normalized to week 0 (left) and normalized to post-stress Week 6 (right) show no significant differences (Mixed ANOVA Week 0 normalization F _(3.705, 27.99)_ = 2.732, p=0.0524, Mixed ANOVA Week 6 normalization F _(2.349, 16.44)_ = 2.730, p= 0.0879). **c**. Susceptible animals trajectory lengths post-event (0-10 sec at onset of CS) during reward trials, normalized to week 0 (left) and normalized to post-stress Week 6 (right) show no significant differences (Mixed ANOVA Week 0 normalization F _(1.006, 5.589)_ = 1.073, p=0.3436; Mixed ANOVA Week 6 normalization F _(0.6656, 3.106)_ = 5.107, p= 0.147). **d**. Control animals trajectory lengths post-event (0-10 sec at onset of CS) during punishment trials, normalized to week 0 (left) and normalized to post-stress Week 6 (right) show no significant differences (Mixed ANOVA Week 0 normalization week F _(3.067, 21.47)_ = 2.169, p=0.1202; Mixed ANOVA Week 6 normalization F _(1.746, 12.22)_ = 1.131, p=0.3462). **e**. Resilient animals trajectory lengths post-event (0-10 sec at onset of CS) during punishment trials, normalized to week 0 (left) were not significantly different across weeks; but when normalized to Post-Stress Week 6 (right) facial feature dynamics show a significant increase in Saline and Post-Ketamine week compared to the Post-Stress baseline (Mixed ANOVA Week 0 normalization week F _(3.264, 24.66)_ = 1.787, p=0.1724; Mixed ANOVA Week 6 normalization F _(1.933, 13.53)_ = 9.185, p=0.0032**, Dunnett’s multiple comparison tests: Week 6 vs. saline p= 0.0121*, Week 6 vs. Ketamine, p=0.3334, Week 6 vs. Post-Ketamine p=0.0008***). **f**. Susceptible animals trajectory lengths post-event (0-10 sec at onset of CS) during punishment trials, normalized to Week 0 (left) and normalized to post-stress Week 6 (right) show no significant differences (Mixed ANOVA Week 0 normalization F _(2.104, 11.69)_ = 1.019, p=0.3947; Mixed ANOVA Week 6 normalization F _(1.371, 6.396)_ = 3.989, p=0.0836).

**Extended Data Figure 13:**
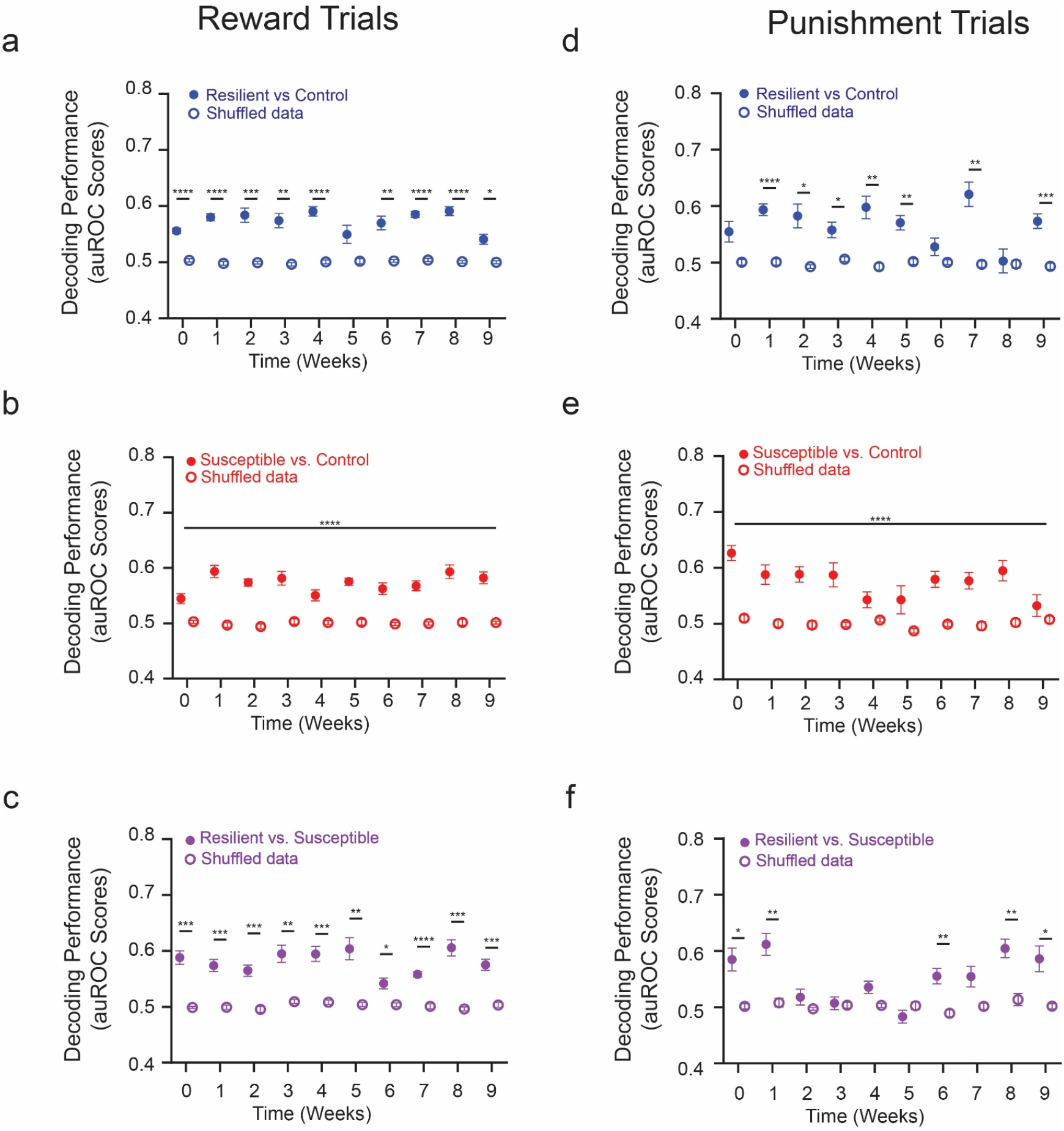
Decoding results for facial dynamics over all weeks. **a**. Decoding performance of resilient vs. control groups compared to shuffled data within individual weeks was significant at all weeks except week 5. Decoding accuracy in response to sucrose in resilient vs. control groups across weeks. (Two-way RM ANOVA week F _(4.618, 83.13)_ = 2.648, p =0.0321; label F _(1,18)_ = 584.6, p<0.0001****; interaction F _(4.618, 83.13)_ = 2.852, p = 0.0229*; Sidak’s multiple comparisons Week 0 p<0.001****, Week 1 p<0.001****, Week 2 p = 0.0008***, Week 3 p = 0.0015**, Week 4 p<0.0001****, Week 5 p = 0.1596, week6 p =0.0027**,Week 7 p<0.0001****, Week 8 p<0.0001****, Week 9 p = 0.0128*). **b**. Decoding performance of *susceptible vs. control* groups was higher overall compared to shuffled data but did not show significant changes across weeks. Decoding accuracy in response to sucrose in *susceptible vs. control* groups across weeks. (Two-way RM ANOVA interaction F _(5.447, 98.05)_ = 2.535, p = 0.0294*, week F _(5.447, 98.05)_ = 2.214, p = 0.0536, label F _(1, 18)_ = 552.4, p<0.0001****). **c**. Significant decoding performance of resilient vs. susceptible groups compared to shuffled data within individual weeks. Decoding accuracy in response to sucrose in resilient vs. susceptible groups across weeks. (Two-way RM ANOVA interaction F _(4.755, 85.60)_ = 2.528, p = 0.0374*, week F _(4.755, 85.60)_ = 2.909, p = 0.0196*, label F _(1, 18)_ = 272.8, p<0.0001****; Sidak’s multiple comparisons Week 0 p =.0004***, Week 1 p = 0.0004***, Week 2 p = 0.0003***, Week 3 p = 0.0028**, Week 4 p = 0.0009***, Week 5 p = 0.0061**, Week 6 p = 0.0330*, Week 7 p<0.0001****, Week 8 p = 0.0002***, Week 9 p = 0.0002***) **d**. Decoding performance of resilient vs. control groups compared to shuffled data within individual weeks was significant at all weeks except weeks 6 and 8. Decoding accuracy in response to air puff in resilient vs. control groups across weeks. (Two-way RM ANOVA interaction F _(4.476, 80.58)_ = 3.881, p = 0.0046**, week F _(4.476, 80.58)_ = 3.361, p = 0.0106*, label F _(1, 18)_ = 229.6, p<0.0001****; Sidak’s multiple comparisons Week 0 p = 0.1506, Week 1 p<0.0001****, Week 2 p = 0.0201*, Week 3 p = 0.0465*, Week 4 p = 0.0038**, Week 5 p = 0.0034**, Week 6 p = 0.7163, Week 7 p = 0.0023**, Week 8 p>0.9999, Week 9 p = 0.0009***). **e**. Significant decoding performance of *susceptible vs. control* groups compared to shuffled data within individual weeks. Decoding accuracy in response to air puff in *susceptible vs. control* groups across weeks. (Two-way RM ANOVA week F _(5.118, 92.12)_ = 2.663, p = 0.0261, label F _(1, 18)_ = 260.5, p<0.0001****, interaction F _(5.118, 92.12)_ = 2.218, p = 0.0576). **f** Significant decoding performance of resilient vs. susceptible groups compared to shuffled data within individual weeks. Decoding accuracy in response to air puff in resilient vs. susceptible groups across weeks. (Two-way RM ANOVA interaction F _(4.899, 88.18)_ = 5.363, p = 0.0003***, week F _(4.899, 88.18)_ = 6.701, p<0.0001****, label F _(1, 18)_ = 120.3, p<0.0001****; Sidak’s multiple comparisons Week 0 p = 0.0253*, Week 1 p =0.0041**, Week 6 p= 0.0076**, Week 8 p= 0.0028**, Week 9 p = 0.0466*)

## References

1. Diagnostic and statistical manual of mental disorders: DSM-5TM, 5th ed. xliv, 947 (American Psychiatric Publishing, Inc., 2013). doi: 10.1176/appi.books.9780890425596

2. Kennedy, S. H. Core symptoms of major depressive disorder: relevance to diagnosis and treatment. Dialogues Clin. Neurosci. 10, 271–277 (2008).

3. Krishnan, V., Han, M.-H., Graham, D. L., Berton, O., Renthal, W., Russo, S. J., Laplant, Q., Graham, A., Lutter, M., Lagace, D. C., Ghose, S., Reister, R., Tannous, P., Green, T. A., Neve, R. L., Chakravarty, S., Kumar, A., Eisch, A. J., Self, D. W., Lee, F. S., Tamminga, C. A., Cooper, D. C., Gershenfeld, H. K. & Nestler, E. J. Molecular adaptations underlying susceptibility and resistance to social defeat in brain reward regions. Cell 131, 391–404 (2007).

4. Ferenczi, E. A., Zalocusky, K. A., Liston, C., Grosenick, L., Warden, M. R., Amatya, D., Katovich, K., Mehta, H., Patenaude, B., Ramakrishnan, C., Kalanithi, P., Etkin, A., Knutson, B., Glover, G. H. & Deisseroth, K. Prefrontal cortical regulation of brainwide circuit dynamics and reward-related behavior. Science 351, aac9698 (2016).

5. Bechara, A., Damasio, H. & Damasio, A. R. Emotion, decision making and the orbitofrontal cortex. Cereb. Cortex N. Y. N 1991 10, 295–307 (2000).

6. Miller, E. K. & Buschman, T. J. Cortical circuits for the control of attention. Curr. Opin. Neurobiol. 23, 216–222 (2013).

7. Euston, D. R., Gruber, A. J. & McNaughton, B. L. The role of medial prefrontal cortex in memory and decision making. Neuron 76, 1057–1070 (2012).

8. Sawaguchi, T. & Goldman-Rakic, P. S. D1 dopamine receptors in prefrontal cortex: involvement in working memory. Science 251, 947–950 (1991).

9. Katz, R. J. Animal model of depression: Pharmacological sensitivity of a hedonic deficit. Pharmacol. Biochem. Behav. 16, 965–968 (1982).

10. Willner, P., Muscat, R. & Papp, M. Chronic mild stress-induced anhedonia: a realistic animal model of depression. Neurosci. Biobehav. Rev. 16, 525–534 (1992).

11. Tye, K. M., Mirzabekov, J. J., Warden, M. R., Ferenczi, E. A., Tsai, H.-C., Finkelstein, J., Kim, S.-Y., Adhikari, A., Thompson, K. R., Andalman, A. S., Gunaydin, L. A., Witten, I. B. & Deisseroth, K. Dopamine neurons modulate neural encoding and expression of depression-related behaviour. Nature 493, 537–541 (2013).

12. Pizzagalli, D. A. Depression, Stress, and Anhedonia: Toward a Synthesis and Integrated Model. Annu. Rev. Clin. Psychol. 10, 393–423 (2014).

13. Destoop, M., Morrens, M., Coppens, V. & Dom, G. Addiction, Anhedonia, and Comorbid Mood Disorder. A Narrative Review. Front. Psychiatry 10, 311 (2019).

14. Whitton, A. E. & Pizzagalli, D. A. Anhedonia in Depression and Bipolar Disorder. Curr. Top. Behav. Neurosci. 58, 111–127 (2022).

15. Lee, J. S., Jung, S., Park, I. H. & Kim, J.-J. Neural Basis of Anhedonia and Amotivation in Patients with Schizophrenia: The role of Reward System. Curr. Neuropharmacol. 13, 750–759 (2015).

16. Dimick, M. K., Hird, M. A., Fiksenbaum, L. M., Mitchell, R. H. B. & Goldstein, B. I. Severe anhedonia among adolescents with bipolar disorder is common and associated with increased psychiatric symptom burden. J. Psychiatr. Res. 134, 200–207 (2021).

17. Forbes, C. E. & Grafman, J. The role of the human prefrontal cortex in social cognition and moral judgment. Annu. Rev. Neurosci. 33, 299–324 (2010).

18. Etkin, A., Egner, T. & Kalisch, R. Emotional processing in anterior cingulate and medial prefrontal cortex. Trends Cogn. Sci. 15, 85–93 (2011).

19. Coley, A. A., Padilla-Coreano, N., Patel, R. & Tye, K. M. Valence processing in the PFC: Reconciling circuit-level and systems-level views. Int. Rev. Neurobiol. 158, 171–212 (2021).

20. Tye, K. M. Neural Circuit Motifs in Valence Processing. Neuron 100, 436–452 (2018).

21. Moda-Sava, R. N., Murdock, M. H., Parekh, P. K., Fetcho, R. N., Huang, B. S., Huynh, T. N., Witztum, J., Shaver, D. C., Rosenthal, D. L., Alway, E. J., Lopez, K., Meng, Y., Nellissen, L., Grosenick, L., Milner, T. A., Deisseroth, K., Bito, H., Kasai, H. & Liston, C. Sustained rescue of prefrontal circuit dysfunction by antidepressant-induced spine formation. Science 364, eaat8078 (2019).

22. Ghosal, S., Duman, C. H., Liu, R.-J., Wu, M., Terwilliger, R., Girgenti, M. J., Wohleb, E., Fogaca, M. V., Teichman, E. M., Hare, B. & Duman, R. S. Ketamine rapidly reverses stress-induced impairments in GABAergic transmission in the prefrontal cortex in male rodents. Neurobiol. Dis. 134, 104669 (2020).

23. Ali, F., Gerhard, D. M., Sweasy, K., Pothula, S., Pittenger, C., Duman, R. S. & Kwan, A. C. Ketamine disinhibits dendrites and enhances calcium signals in prefrontal dendritic spines. Nat. Commun. 11, 72 (2020).

24. Drysdale, A. T., Grosenick, L., Downar, J., Dunlop, K., Mansouri, F., Meng, Y., Fetcho, R. N., Zebley, B., Oathes, D. J., Etkin, A., Schatzberg, A. F., Sudheimer, K., Keller, J., Mayberg, H. S., Gunning, F. M., Alexopoulos, G. S., Fox, M. D., Pascual-Leone, A., Voss, H. U., Casey, B., Dubin, M. J. & Liston, C. Resting-state connectivity biomarkers define neurophysiological subtypes of depression. Nat. Med. 23, 28–38 (2017).

25. Mayberg, H. S., Lozano, A. M., Voon, V., McNeely, H. E., Seminowicz, D., Hamani, C., Schwalb, J. M. & Kennedy, S. H. Deep Brain Stimulation for Treatment-Resistant Depression. Neuron 45, 651–660 (2005).

26. Dolensek, N., Gehrlach, D. A., Klein, A. S. & Gogolla, N. Facial expressions of emotion states and their neuronal correlates in mice. Science 368, 89–94 (2020).

27. Moëne, O. L. & Larsson, M. A New Tool for Quantifying Mouse Facial Expressions. eNeuro 10, (2023).

28. Cunningham, J. P. & Yu, B. M. Dimensionality reduction for large-scale neural recordings. Nat. Neurosci. 17, 1500–1509 (2014).

29. Pachitariu, M., Stringer, C., Dipoppa, M., Schröder, S., Rossi, L. F., Dalgleish, H., Carandini, M. & Harris, K. D. Suite2p: beyond 10,000 neurons with standard two-photon microscopy. 061507 Preprint at 10.1101/061507 (2017)

30. Padilla-Coreano, N., Batra, K., Patarino, M., Chen, Z., Rock, R. R., Zhang, R., Hausmann, S. B., Weddington, J. C., Patel, R., Zhang, Y. E., Fang, H.-S., Mishra, S., LeDuke, D. O., Revanna, J., Li, H., Borio, M., Pamintuan, R., Bal, A., Keyes, L. R., Libster, A., Wichmann, R., Mills, F., Taschbach, F. H., Matthews, G. A., Curley, J. P., Fiete, I. R., Lu, C. & Tye, K. M. Cortical ensembles orchestrate social competition through hypothalamic outputs. Nature 603, 667–671 (2022).

31. Pereira, T. D., Tabris, N., Matsliah, A., Turner, D. M., Li, J., Ravindranath, S., Papadoyannis, E. S., Normand, E., Deutsch, D. S., Wang, Z. Y., McKenzie-Smith, G. C., Mitelut, C. C., Castro, M. D., D’Uva, J., Kislin, M., Sanes, D. H., Kocher, S. D., Wang, S. S.-H., Falkner, A. L., Shaevitz, J. W. & Murthy, M. SLEAP: A deep learning system for multi-animal pose tracking. Nat. Methods 19, 486–495 (2022).

